# Type VI collagen is proportionally lower around airways and blood vessels in idiopathic pulmonary fibrosis

**DOI:** 10.64898/2026.01.13.698558

**Authors:** Helene Wallem Breisnes, Mehmet Nizamoglu, Maunick Lefin Koloko Ngassie, Theo Borghuis, Marnix Roel Jonker, Rhode Meuleman, Tessa Kole, Corry-Anke Brandsma, Simon Francis Thomsen, Morten Karsdal, Diana Julie Leeming, Filipa Bica Simões, Jannie Marie Bülow Sand, Janette Kay Burgess

## Abstract

Type VI collagen (COL6) is a key extracellular matrix protein that supports matrix organization and cell-matrix interactions, yet its regulation in idiopathic pulmonary fibrosis (IPF) remains poorly understood. Here, we characterize COL6 gene expression, spatial localization, remodeling, and functional effects of COL6-derived fragments. Analysis of publicly available single-cell RNA sequencing data from 30 controls and 32 pulmonary fibrosis patients revealed higher expression of *COL6A1–A6* in fibrotic lungs, predominantly in mesenchymal cells (*A1*: p=0.0002, *A2*: p=0.0005, *A3*: p=2.5x10^-5^, *A5*: p=0.016, *A6*: p=0.007). Immunohistochemical analysis of lung tissue from never-smoker (n=9), ex-smoker (n=9) controls, and IPF patients (n=12) showed extensive COL6 localization across parenchyma, airways, and vessels. The proportion of COL6α1 and COL6α2 was lower around IPF vessels (α1: p<0.001, α2: p=0.012), and COL6α2 was lower around IPF airways (p=0.033) compared with never-smokers. Quantification of COL6 remodeling fragments in lung tissue from never-smoker (n=3), ex-smoker (n=5) controls, and IPF patients (n=10) revealed that COL6 production (PRO-C6) localized around airways and vessels but was proportionally lower in IPF airways (never-smokers: p=0.0075). In contrast, COL6 degradation (C6M) was widely distributed throughout the tissue, with lower levels in IPF (never-smokers: p=0.0008). Functionally, COL6 and PRO-C6 increased fibroblast viability (COL6: p=0.002, PRO-C6: p=0.0021), while apoptosis was unaffected. Similar trends were observed in epithelial and endothelial cells. In summary, despite increased COL6 gene expression, IPF lungs exhibited lower proportions of COL6 protein and synthesis around airways and vessels, suggesting disrupted matrix organization, altered remodeling, and pro-survival effects that may contribute to fibroblast persistence and fibrosis progression.

**New & Noteworthy:** In this study, we revealed that IPF lungs are characterized by increased type VI collagen (COL6) gene expression but lower COL6 protein and turnover proportions around airways and blood vessels compared with controls. COL6 and a fragment associated with its production (PRO-C6) increased lung fibroblast viability. These findings highlight that pulmonary fibrotic tissue is characterized by disrupted extracellular matrix and altered tissue remodeling and suggest that COL6 may contribute to fibroblast persistence and fibrosis progression.

## Introduction

Idiopathic pulmonary fibrosis (IPF) is a chronic and rapidly progressive interstitial lung disease, characterized by fibrotic scarring of the lung tissue. In healthy lungs, there is continuous extracellular matrix (ECM) remodeling that sustains tissue homeostasis. In contrast, in IPF the balance between ECM production and degradation is disrupted, resulting in excessive ECM accumulation, disruption in the component proportions within the ECM, and distortion of the lung architecture [1–4].

Type VI collagen (COL6) is a unique and essential component of the ECM, located at the interface between the basement membrane (BM) and interstitial matrix (IM) [5]. In the lung, it is found near airways, alveolar epithelium, and vascular endothelium [6]. Of the 28 identified collagens, COL6 is recognized as one of the most abundant in the mammalian lung, where it orchestrates the organization of the tissue architecture and facilitates cell attachment and linkage to the surrounding matrix [7–9]. There are six known α chains of this collagen, encoded by the six genes *COL6A1-A6.* While all six genes exist in humans, *COL6A4* is nonfunctional due to a chromosome inversion, rendering the α4 chain absent in humans [10]. The α1, α2, and α3 are the primary chains forming the traditional collagen triple-helix structure, whereas α5 and α6 are homologues of α3 and have been suggested as potential substitutes with specific tissue localizations [9–11]. In IPF, COL6 gene expression has been shown to be upregulated compared with controls [12,13] with no differential regulation between the α1 and α3 chains [13].

Various cell types produce COL6; however, fibroblasts are considered the primary source [14]. Production and maturation of COL6 occurs intracellularly, where two monomers associate in an antiparallel manner, followed by formation of tetramers in a parallel manner [15]. Upon secretion of mature tetramers from a cell to facilitate incorporation of COL6 in the ECM, the C5 domain of the α3 chain is cleaved off. This process releases a bioactive fragment – endotrophin – that can be measured through detection of the resulting neoepitope that is exposed during the cleavage process. COL6 tetramers assemble to form a unique beaded-filament structure that then aggregates to form a microfibrillar network within the deposited ECM in the tissue [16]. To maintain tissue homeostasis, matrix metalloproteinases (MMPs) mediate tissue degradation, which results in the release of ECM degradation fragments into circulation, where they can be measured as serological biomarkers. Notably, several studies have found increased circulating levels of neoepitopes reflecting COL6 production and endotrophin (PRO-C6) and MMP-mediated degradation (C6M) in patients with IPF, which were associated with disease severity, progression, and mortality [17–20]. These findings are also mirrored in other fibrotic conditions, such as skin, kidney, and liver [21–24]. An overview of the COL6 composition, maturation, and remodeling is shown in **Figure 1**.

**Figure 1.**
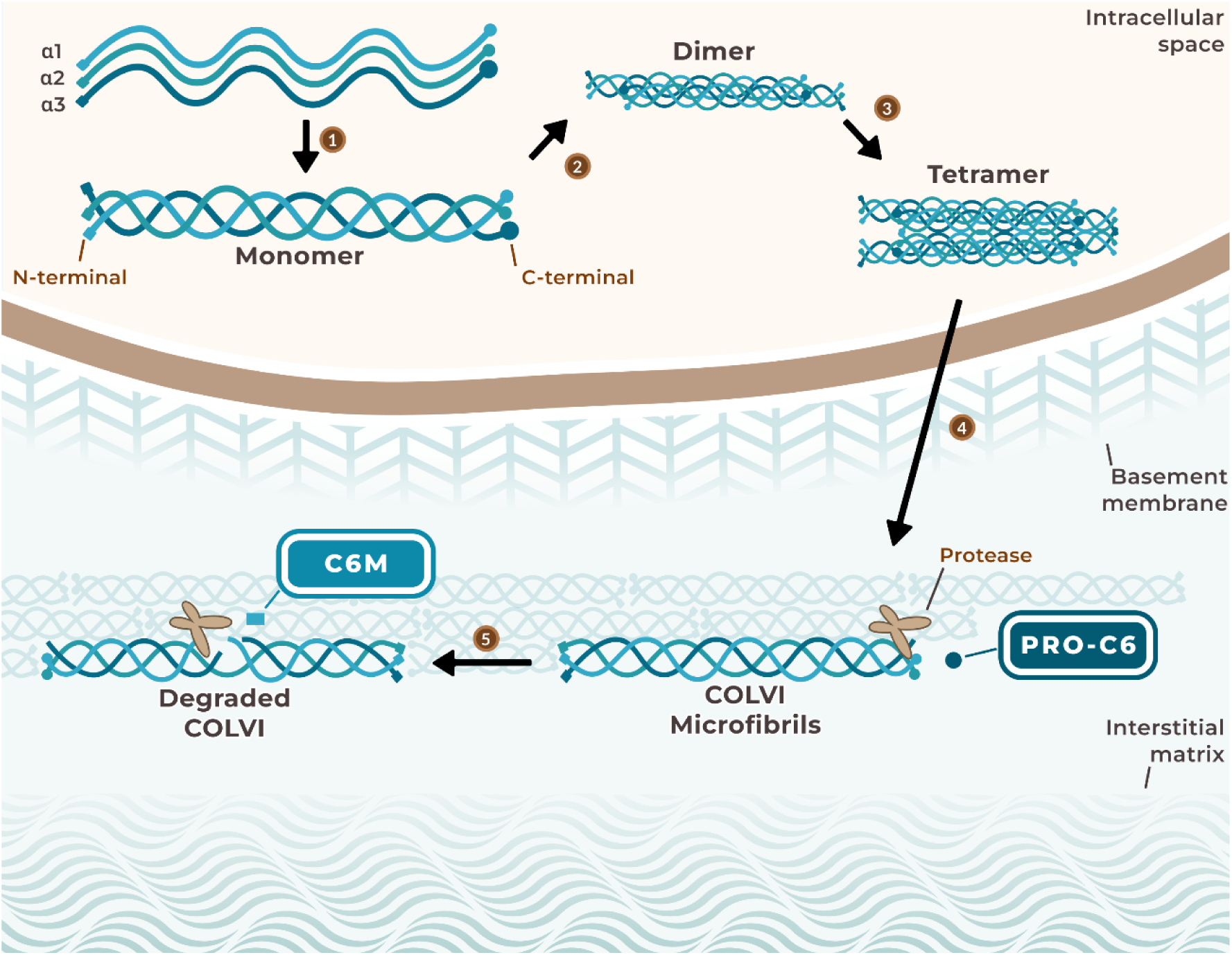
COL6 composition, maturation, and remodeling in the lung ECM. COL6 is composed of three main α chains (α1-α3) with N- and C-terminal globular domains, (1) which assemble into a traditional collagen triple-helical monomer. (2) Intracellularly, two monomers associate in an antiparallel manner, (3) followed by the formation of tetramers in a parallel orientiation. (4) These tetramers are secreted and assembled into a unique microfibrillar network at the interface between the basement membrane and interstitial matrix. During incorporation into the ECM, the C5 domain at the C-terminal end of the α3 chain (PRO-C6) is cleaved off. (5) During tissue degradation, matrix metalloproteases cleave COL6, releasing degradation fragments (C6M). COL6/COLVI: type VI collagen, ECM: extracellular matrix.

COL6 supports cell survival, as illustrated in mouse fibroblasts [25] and megakaryocytes [26]. However, recent research has focused on the role of its bioactive fragment, endotrophin. Endotrophin is recognized for its signaling properties and pro-fibrotic functions which have been associated with increased expression of ECM regulating genes [27–29]. More specifically, endotrophin can act as a co-stimulator of pathological pathways leading to fibrosis and inflammation in adipose tissue [29]. In another study, urinary endotrophin was associated with T cell-mediated rejection in kidney transplant recipients, raising the question whether endotrophin modulates T cell response in kidneys [30]. These recent studies highlight how COL6 and its associated fragments can influence the cell and tissue microenvironment in both health and disease.

Despite the abundance of COL6 in the lungs, research describing COL6 remodeling and released fragments associated with this remodeling during pulmonary fibrosis remains sparse. Thus, in this study we aimed to explore COL6 gene and protein expression, COL6 remodeling through its production and degradation fragments, together with their influences on different lung resident cell types.

## Materials and Methods

### Experimental Design

This study investigated the role of COL6 in fibrotic lung disease by combining transcriptomic, protein, and functional analyses using human lung tissue from pulmonary fibrosis patients and cell cultures from non-diseased lung cells. A schematic overview of the experimental workflow and methodological details is presented in **Figure 2**.

**Figure 2.**
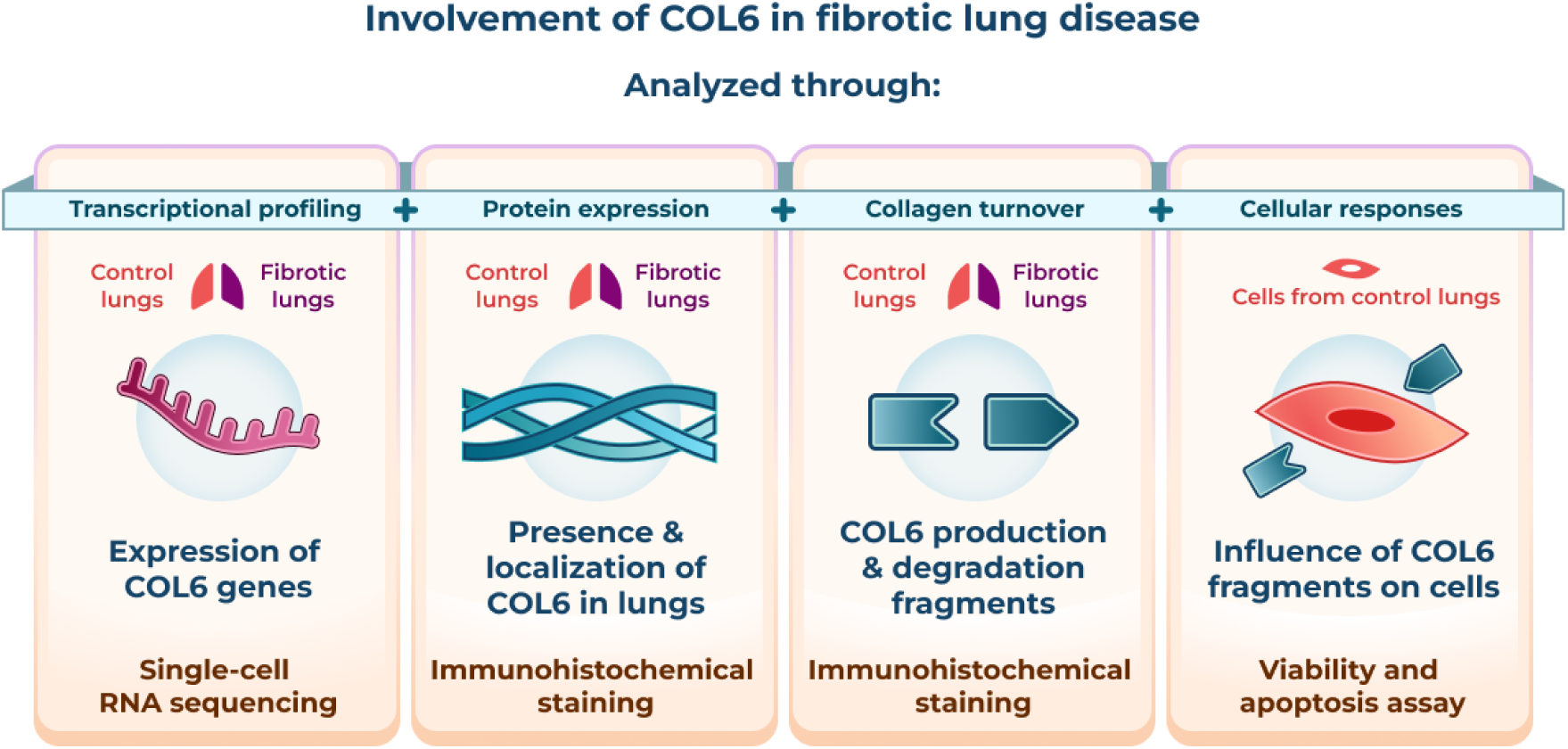
Overview of the experimental approach and analysis strategy employed in this study. The involvement of COL6 was investigated through its gene expression, protein expression and turnover, and effects on cell viability and apoptosis. Gene expression was analyzed using single-cell RNA sequencing publicly available data from fibrotic and control lungs. The localization of COL6 chains and production (PRO-C6) and degradation (C6M) fragments were detected using IHC staining on control and IPF lung tissue samples with digital image analysis. Lastly, in vitro cell-based experiments were conducted to assess the influence of COL6 whole protein, PRO-C6, and C6M on cell viability and apoptosis. C6M: MMP-mediated type VI collagen degradation, COL6: type VI collagen, IHC: immunohistochemistry, PRO-C6: type VI collagen production.

### Single-cell RNA sequencing data analysis

Gene expression profiles of genes encoding different COL6 chains (*COL6A1*, *COL6A2*, *COL6A3*, *COL6A5,* and *COL6A6*), which encode the α1-3, α5, and α6 chains, were analyzed across major cell populations in lung tissue of patients with lung fibrosis and control donors using a publicly available single-cell RNA sequencing (scRNAseq) dataset generated by Mayr *et al.* [31]. The dataset includes samples from control donors (n=30) and patients with fibrosis (n=32) collected from three different cohorts: the Chicago cohort (<u>GSE122960</u>) [32], the Nashville cohort (<u>GSE135893</u>) [33], and the Munich cohort [34]. The integrated dataset was re-analyzed using Scanpy version 1.10.4 in Python version 3.13.1 [35], without further modifications to the original data. Four major cell populations annotated by Mayr et al. were used: CLND5^+^ endothelial cells, EPCAM^+^ epithelial cells, PTPRC^+^ immune cells, and COL1A2^+^ mesenchymal cells. Gene expression levels were quantified using a pseudobulk approach, where raw single-cell counts were aggregated by donor and cell type to generate sample-level expression profiles. Plots for the statistical comparisons were generated using the Python packages matplotlib version 3.10.0 and seaborn version 0.13.2.

### Lung tissue collection

The lung tissue samples used in this study were obtained from left-over lung material from tumor resection (non-diseased controls) or lung transplantation (IPF) surgeries at the University Medical Center Groningen (UMCG). Control tissue used for immunohistochemical staining and fibroblast isolation was collected from areas as distant as possible from the tumor and exhibited a macroscopically normal appearance as confirmed by a pathologist.

Epithelial cells were collected from endobronchial brushes obtained during bronchoscopy in healthy control individuals recruited for the “ARMS cohort (NCT03141814, “Asthma Origins and Remission Study”) at the UMCG. The ARMS study was approved by the Medical Ethical Committee and conducted in accordance with the research code and the national ethical and professional guidelines “Code of conduct for Health Research”.

This study was conducted in compliance with the Research Code of the University Medical Center Groningen (UMCG), as outlined in https://umcgresearch.org/w/research-code-umcg, and adhered to national ethical and professional guidelines Code of Conduct for Health Research (https://www.coreon.org/wp-content/uploads/2023/06/Code-of-Conduct-for-Health-Research-2022.pdf). The use of data and leftover lung tissue in this study was exempt from the Medical Research Human Subjects Act in The Netherlands, as confirmed by the Medical Ethical Committee of the UMCG. The study protocol was approved by the Central Ethics Review Board non-WMO studies (study no. 10748) and under Dutch laws (Medical Treatment Agreement Act [WGBO] art. 458; GDPR art. 9; UAVG art. 24) was exempt from the requirement for informed consent. All donor material and clinical data were deidentified before experimental use, ensuring investigators had no access to identifiable information.

### Immunohistochemical staining

The relative amounts of COL6α1 and COL6α2 were quantified through immunohistochemical (IHC) staining. An overview of patient characteristics for donors of control lung tissue (n=9 for both never-smoker and ex-smoker) and IPF lung tissue (n=12) is provided in **Table 1**. The tissue samples were part of the HOLLAND (HistopathOLogy of Lung Aging aNd COPD) project and were processed as described previously [36–39]. Briefly, samples were fixed in formalin, embedded in paraffin, and sectioned at 6 μm thickness. Serial sections were stained for COL6α1 and COL6α2 using primary antibodies diluted 1:3200 and 1:12500, respectively (COL6α1: polyclonal rabbit, anti-human, cat. no. NB120-6588, Novus Biologicals, Centennial, CO, USA and COL6α2: monoclonal rabbit, anti-human, cat. no. ab180855, Abcam, Cambridge, UK). The secondary antibody was a polyclonal goat anti-rabbit antibody (cat. no. P0488, Dako, Denmark) for both targets. Positively-stained target proteins were visualized using Vector NovaRED Substrate (cat. no. SK-4800, Vector Laboratories, Newark, USA). Sections were counterstained using hematoxylin. Tissue slides were imaged using a Hamamatsu NanoZoomer digital scanner (Hamamatsu Photonic K.K., Shizuoka, Japan), as described previously [39].

**Table 1.** Patient characteristics of tissue donors for COL6α1 and COL6α2 IHC staining.

|  | Never-smoker controls | Ex-smoker controls | IPF | Never-smoker controls vs IPF | Ex-smoker controls vs IPF |
| --- | --- | --- | --- | --- | --- |
| **n** | 9 | 9 | 12 |  |  |
| **Age (years), median (range)** | 70 (50-82) | 55 (43-62) | 58.50 (37-67) | 0.0213* | ns* |
| **Male sex, n (%)** | 1 (11) | 3 (33) | 9 (75) | 0.075# | ns# |
| **FEV1 (%pred), median (range) †** | 107.5 (80-121) | 103.6 (85-127) | 49.35 (18-74.5) | 0.0002* | 0.0002* |

Data are shown as median with IQR or as counts (n, %). Groups were compared by \*: Mann-Whitney test (for continuous variables) or #: Fisher's exact test (for categorical variables). †: Data available for n=8 across all groups. IHC: immunohistochemistry, IPF: Idiopathic pulmonary fibrosis, FEV1 (%pred): % Predicted Forced expiratory volume in 1 second. ns: not significant.

Staining against neoepitopes of COL6 production (nordicPRO-C6^TM^) [40] and MMP-mediated degradation (nordicC6M^TM^) [41] was performed as described in the previous section unless otherwise stated. The following neoepitope-specific primary antibodies were diluted 1:2000 and 1:100, respectively, in 2% BSA-PBS (PRO-C6: monoclonal mouse, anti-human, cat. no. 4000AG01, Nordic Bioscience, Herlev, Denmark and C6M: monoclonal mouse, anti-human, cat. no. 1500AG01, 1:100, Nordic Bioscience). The tissue slides were incubated with the primary antibodies for 1 h at room temperature. Absorption controls were performed by preincubating the primary antibody with a synthetic peptide corresponding to its recognized epitope (standard peptides provided by the producer of antibodies, Nordic Bioscience) for 1 h at room temperature at a 1:3 ratio. An overview of patient characteristics of the tissue donors stained for COL6 fragments is provided in **Table 2**. Notably, one of the never-smoker controls and one of the patients with IPF were the same between the two cohorts.

**Table 2.**
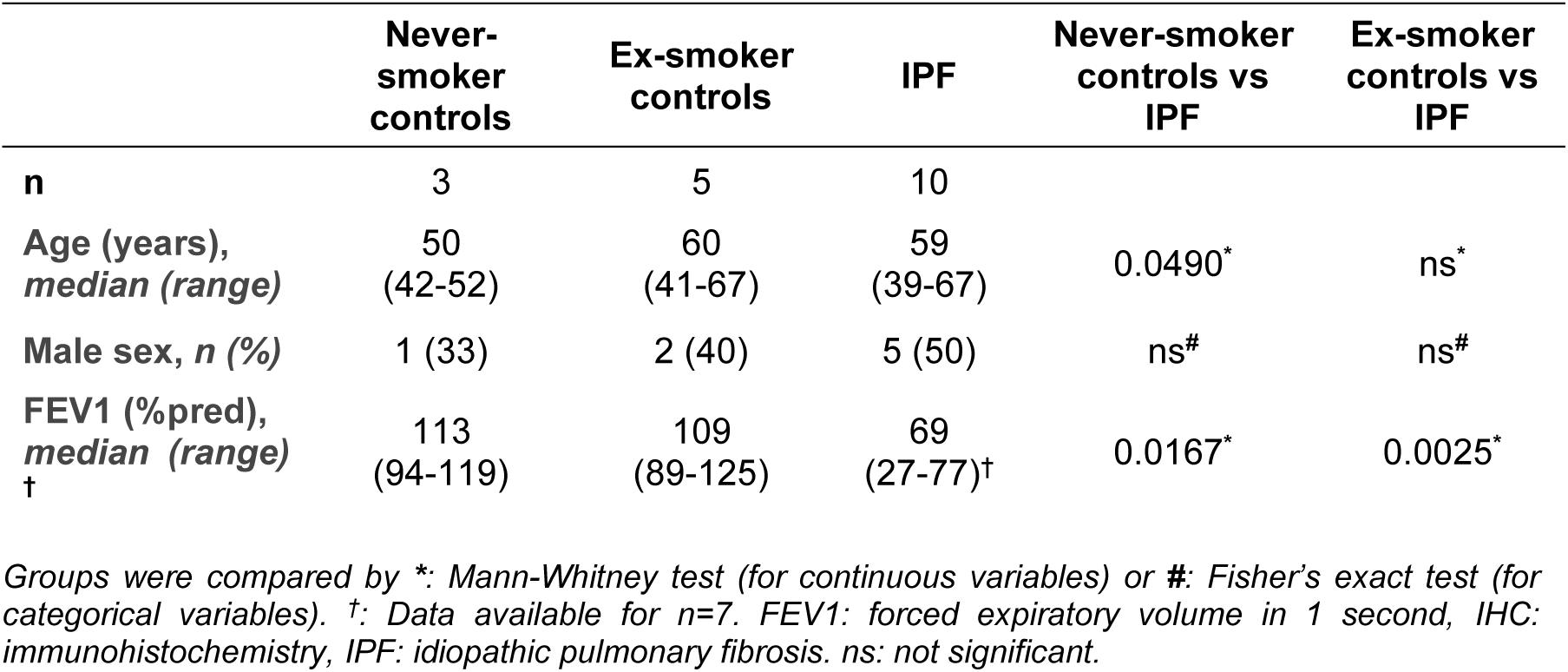
Patient characteristics of tissue donors for PRO-C6 and C6M IHC staining.

### Image analysis

Scanned images of lung-tissue sections stained for COL6α1, COL6α2, PRO-C6, and C6M from controls and patients with IPF were analyzed as previously described [39]. Whole-tissue images were processed following artifact removal using Adobe Photoshop^®^ version 25.9.1 (Adobe Inc., CA, USA). Parenchyma, airway walls, and blood vessel regions were extracted from the whole-tissue images using Aperio ImageScope version 12.4.6 (Leica Biosystems, Germany). Regions corresponding to IPF airway walls were identified according to our previously described method [38].

The total tissue areas positive for the target proteins were quantified by applying color deconvolution to separate the different staining colors. Fiji/Image J version 1.54j [42] was used for automatic image processing, as previously described [39]. Subsequently, RStudio version 2024.04.2 was used to sort the raw data, and the percentage of positively-stained tissue area was calculated using the following formula.

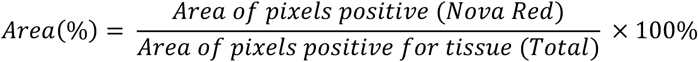

### Cell isolation and maintenance

Non-diseased control primary human lung fibroblasts (n=5) were isolated and frozen for storage, following previously described protocols [43]. The patient characteristics of the donors are summarized in **Table 3**. Frozen fibroblasts were retrieved from liquid nitrogen maintained in complete growth medium consisting of Dulbecco’s Modified Eagle Medium (DMEM low glucose [1 g/L]; Gibco, Thermo Fisher Scientific, Waltham, MA, USA) supplemented with 10% fetal bovine serum (FBS; Sigma Aldrich, St. Louis, MO, USA) and 1% Penicillin-Streptomycin (P/S; Gibco).

**Table 3.**
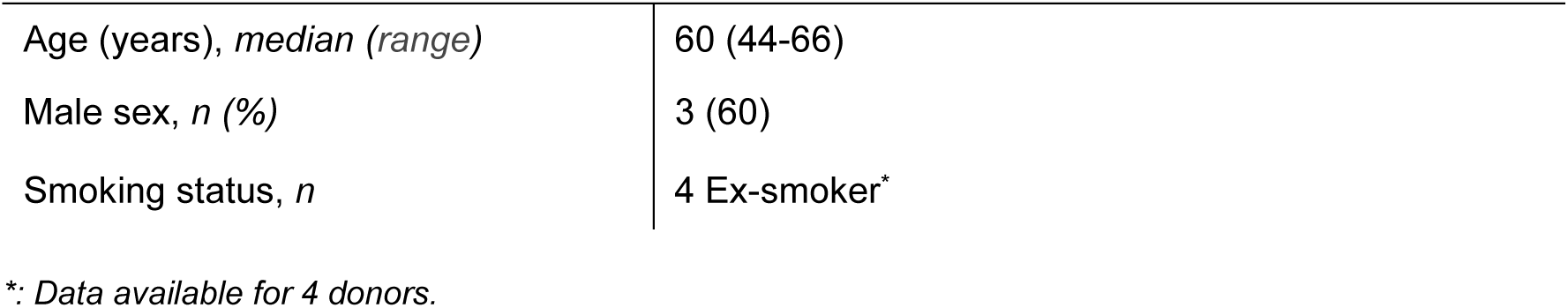
Patient characteristics of the fibroblast donors (n=5).

Once confluent, fibroblasts were harvested using 0.25% trypsin-EDTA (Gibco) and centrifuged at 500 g for 5 min. Afterwards, supernatants were discarded and cell pellets were resuspended in 1 mL complete growth medium and counted using a Bürker-Turk counting chamber. 20000 cells per well were seeded in a 96-well plate and incubated for 24 h at 37°C in 5% carbon dioxide (CO_2_). Fibroblasts were quiesced by replacing the growth medium with quiescence medium (DMEM low glucose [1 g/L] supplemented with 1% P/S) and further incubated for 24 h at 37°C and 5% CO_2_. After this period, treatments were applied to investigate effects on cell viability and apoptosis.

Brushed epithelial cells (n=3) from healthy control individuals were isolated using previously described protocols [44]. Donor characteristics are summarized in **Table 4**. The isolated epithelial cells were thawed from liquid nitrogen storage and maintained in Airway Epithelial Growth Medium (AEGM; PromoCell, Heidelberg, Germany) enriched with the accompanied BulletKit and 1% P/S. Once confluent, cells were detached using 0.25% trypsin-EDTA and upscaled for one additional passage. Cells at passage four were harvested and counted using a Bürker-Turk counting chamber. A total of 20000 cells per well were seeded in 96-well plates and cultured for 24 h at 37°C and 5% CO_2_, followed by overnight quiescence in AEGM supplemented with 10 µg/mL transferrin (Sigma Aldrich), 5 µg/mL Insulin (Sigma Aldrich), and P/S. The next morning, cells were stimulated with treatments.

**Table 4.** Patient characteristics of the epithelial cell donors (n=3).

|  |  |
| --- | --- |
| Age (years), *median (range)* | 54 (39-65) |
| Male sex, *n (%)* | 2 (66) |
| Smoking status, *n* | 1 Ex-smoker* |

\*: Data available for 2 donors.

Human pulmonary microvascular endothelial cells (HPMEC-ST1.6R) were a kind gift from Dr. Kirkpatrick (Johannes Gutenberg University, Mainz, Germany) [45]. Cells (n=3) were thawed from liquid nitrogen storage at passage 29 and maintained in MCDB131 medium (Gibco) supplemented with 10% FBS, 1% GlutaMAX, 10 ng/ml endothelial growth factor (homemade bovine brain extract), 1 ug/ml hydrocortisone (Invitrogen, Thermo Fisher Scientific, Waltham, MA, USA) and 1% P/S. Once confluent, cells were detached using 0.25% trypsin-EDTA and upscaled for an additional passage. Cells at passage 31 were harvested and counted using a Bürker-Turk counting chamber in an n=3. A total of 20000 cells per well were seeded in 96-well plates and cultured for 24 h at 37°C and 5% CO_2_, followed by overnight quiescence in MCDB131 medium supplemented with 0.5% FCS and P/S. The following morning, cells were stimulated with treatments.

### Cell treatment and viability and apoptosis assays

To treat the different cells, a final volume of 100 µL of complete growth medium was used per well. For the negative control, cells were treated with growth medium alone, whereas for COL6 (cat. no. ab7538, Abcam), PRO-C6, and C6M (both provided by Nordic Bioscience) a final concentration of 8 ng/μL was used. The concentrations were decided based on a dose response curve. Afterwards, the plate was incubated for 24 h at 37°C in 5% CO_2_, followed by viability and apoptosis assays.

After 24 h of incubation with treatments, reagents were prepared for the viability and apoptosis assay following the ApoLive-Glo Multiplex Assay Quick Protocol for 96-well plates (cat. no. G6410, Promega, Madison, WI, USA). This assay enables assessment of the number of viable cells and activation of caspase 3/7 within a single well. The assay was run according to the official protocol. Subsequently, fluorescence absorbance (viability) was measured at 400 nm for excitation and 505 nm for emission, followed by measurement of luminescence (caspase 3/7 activity), using a ClarioStar Plus Spectrophotometer (BMG Labtech, Ortenberg, Germany).

### Statistical analysis

Differences in gene expression profiles of COL6 α chains between IPF and control groups, generated by pseudobulk step using publicly available scRNAseq data, were analyzed using the Mann-Whitney test. For quantified image analysis, depending on the normal distribution, ordinary one-way ANOVA or Kruskal-Wallis test was used for comparing whole tissue and parenchyma images among different groups and shown as mean ± standard deviation (SD) or median ± interquartile range (IQR), respectively. A linear mixed model was used for comparison of groups to account for multiple airways or blood vessels per patient, including a random intercept for each tissue and an unstructured covariance structure, and data are shown as median ± IQR. Cell viability and apoptosis data were log-transformed and analyzed using a paired t-test. Data are shown as fold change from the negative control. Data were considered statistically significant if *p* < 0.05. Analyses were conducted using R Studio version 2024.04.2 + 764 and GraphPad Prism V.10.2.3.

## Results

### COL6 is mainly expressed in mesenchymal cells and higher in fibrotic lungs

The Mayr *et al*. dataset [31], which includes control and fibrotic lung samples from multiple cohorts, enabled evaluation of COL6 subtype expression across disease states. When these data were compartmentalized into four major cell types (endothelial, epithelial, immune, and mesenchymal cells) (**Figure 3A**), gene expression levels of COL6 α chains were predominantly found in mesenchymal cells (**Figure 3B**). Pseudobulk analysis revealed distinct gene expression patterns for each COL6 α chain between control and fibrotic samples in all major cell types (**Figure 3C**, detailed violin plots are presented in **Supplementary Figure 1**). *COL6A1* and *COL6A3* gene expression were higher in fibrotic endothelial (*A1*: p=0.013, *A3*: p=0.044), immune (*A1*: p=9.2 x 10^-5^, *A3*: p=0.017), and mesenchymal (*A1*: p=1.9 x 10^-4^, *A3*: p=2.5 x 10^-5^) cells compared with controls. *COL6A2* gene expression was significantly higher in all four major cell types in fibrotic lungs (endothelial cells: p=0.003, epithelial cells: p=0.014, mesenchymal cells: p=2.2x10^-4^, immune cells: p=4.9x10^-4^), while *COL6A5* and *COL6A6* were only higher in fibrotic mesenchymal cells (*A5*: p=0.016, *A6*: p=0.007) compared with controls.

**Figure 3.**
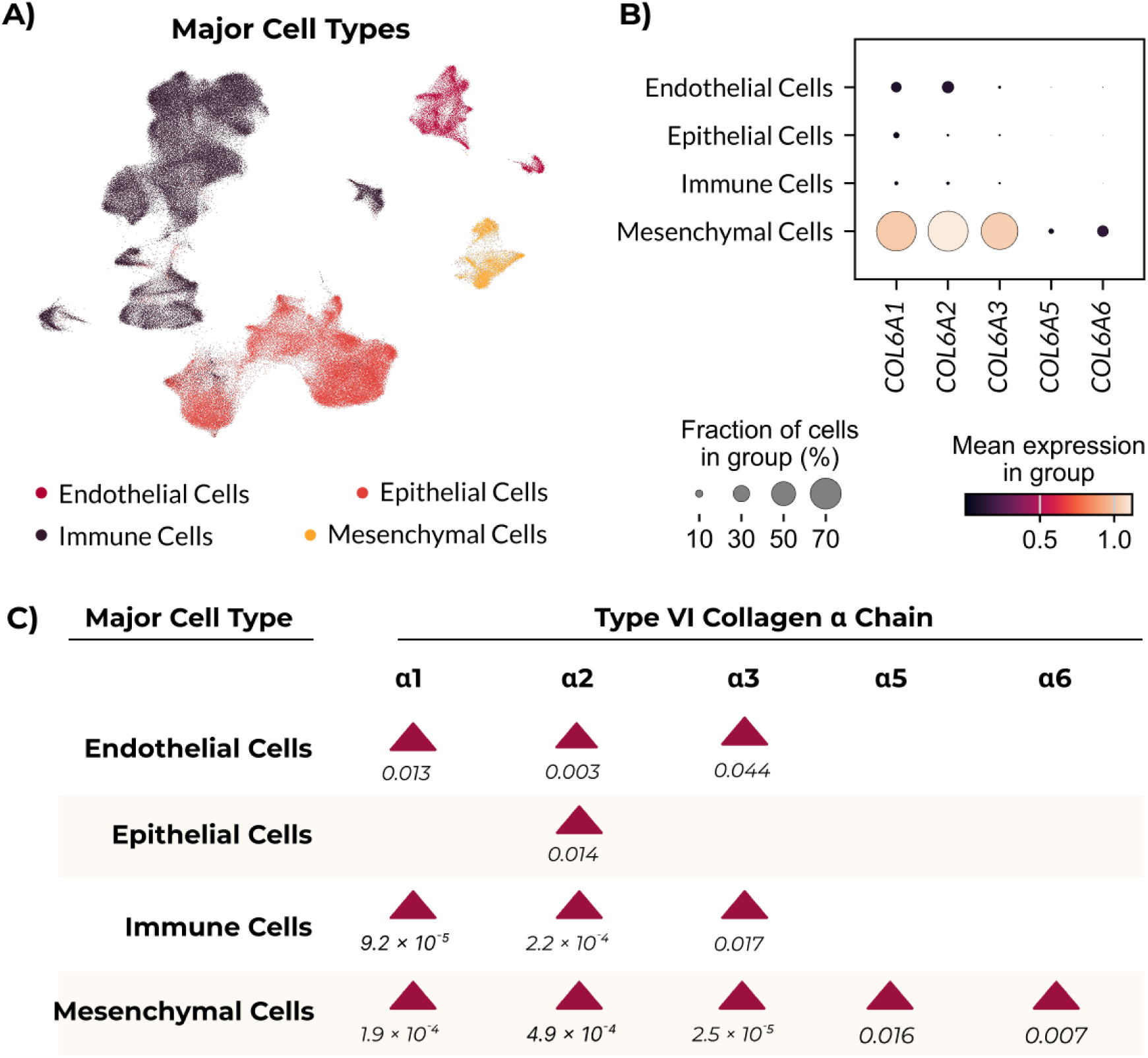
COL6 α chain gene expression levels are higher in cells from fibrotic lung samples compared with controls. Gene expression profiles of control and fibrotic lungs were analyzed in a publicly available single-cell RNA sequencing dataset [31]. **A)** UMAP plot showing the distribution of major cell types within the dataset. **B)** Dot plot depicting the fraction of cells expressing genes encoding the different COL6 α chains. **C)** Pseudobulk analysis comparing COL6 α chains in control (n=30) and fibrotic (n=32) lungs. Red triangles indicate significantly higher gene expression levels of COL6A1-6 (columns) in fibrotic lungs for a given cell type (rows) compared with controls. Statistical comparison for panel **C**: Mann-Whitney test. COL6: type VI collagen.

### Fibrotic airways have proportionally less COL6α2

IHC was performed on human lung tissue sections from never-smoker and ex-smoker controls, as well as patients with IPF, to determine the extent and spatial distribution of COL6α1 and COL6α2.

In controls, COL6α1 primarily localized around the airways and blood vessels, with a less distinct distribution in the parenchyma (**Figure 4A-B)**. In IPF, heterogeneous staining was seen throughout the entire tissue (**Figure 4C**). Whole-tissue image analysis – which was normalized to the total amount of tissue – revealed that a lower proportion of the tissue area was positively stained for COL6α1 in IPF compared with ex-smoker (p=0.0395, **Supplementary Figure 2A**) but not never-smoker controls. Additionally, a lower average staining intensity – across pixels that were positive for COL6α1 – were found in IPF compared with never-smoker and ex-smoker controls (p=0.0003 and p<0.0001, respectively, **Supplementary Figure 2E**). Individual regions within the tissue sections were analyzed using the same approach. In parenchyma, there were no differences in the proportion of COL6α1, but the average staining intensity of positive pixels was lower in IPF compared with never-smoker and ex-smoker controls (p=0.0122 and p=0.0005, respectively, **Supplementary Figure 2B & F**). In the airways, no differences were observed in either the proportion of COL6α1 or the positive pixel average staining intensity (**Supplementary Figure 2C & G**). Around the blood vessels, there was a significantly lower proportion of COL6α1 and lower average positive pixel staining intensity in IPF compared with never-smoker and ex-smoker controls (never-smoker: %area: p<0.001, intensity: p<0.001, ex-smoker: %area: p=0.006, intensity: p<0.001, **Figure 4D-E**). The proportion of COL6α1 was also lower in ex-smokers compared with never-smoker controls (p=0.006, **Supplementary Figure 2D**). In summary, COL6α1 tissue changes were most pronounced around IPF blood vessels where both the protein proportion and average positive pixel staining intensity were lower compared with controls.

**Figure 4.**
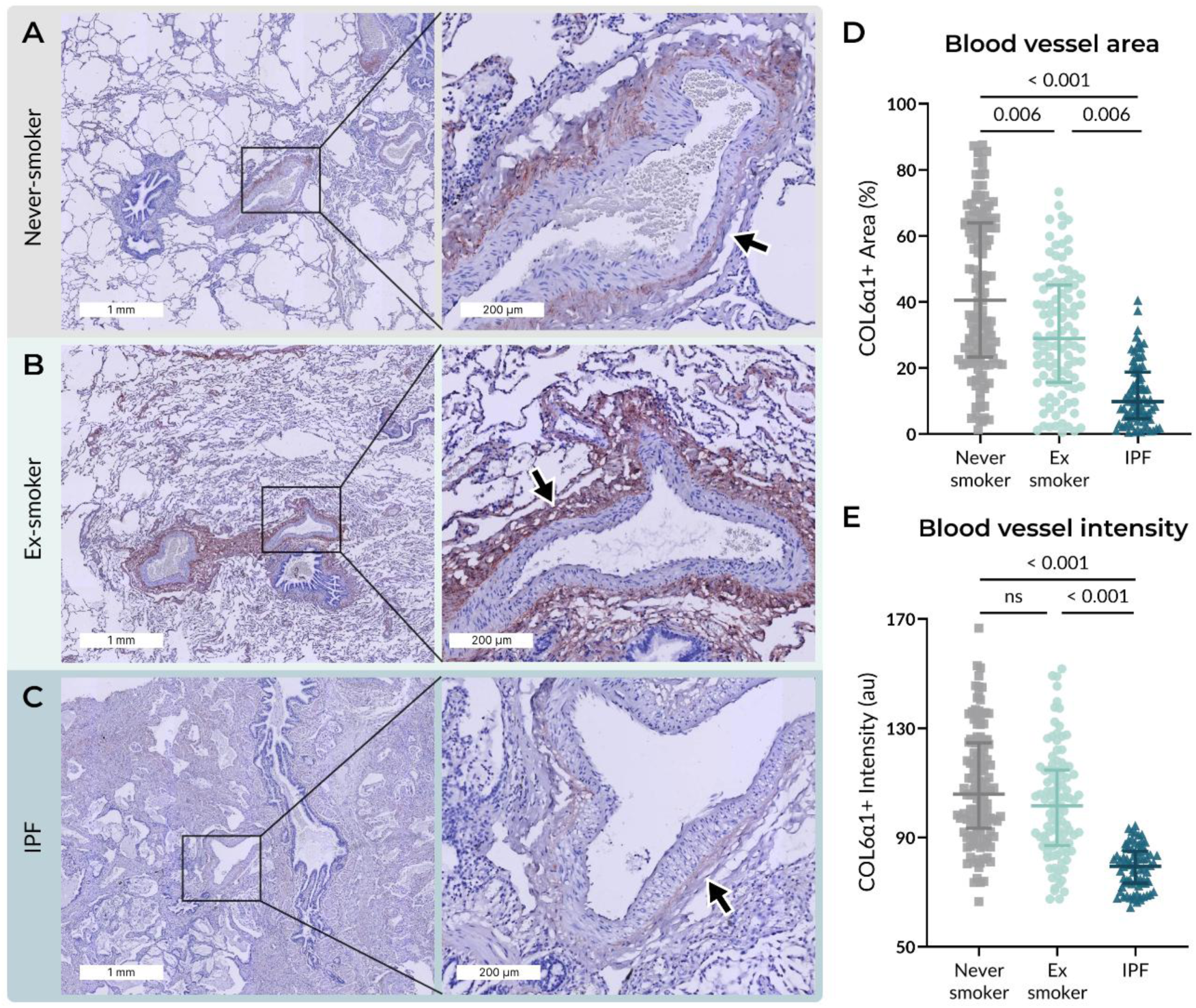
IPF blood vessels have relatively lower COL6α1 compared with controls. Images of sections of lung tissue stained for COL6α1 were digitally quantified. Representative tissue images from **A)** never-smoker control, **B)** ex-smoker control, and **C)** IPF subjects. Scale bars: 1 mm for images on the left; 200 µm for images on the right. The arrows indicate blood vessels. Contrast and saturation were digitally increased by 5% for all images to enhance visibility. **D)** Positively-stained area (%) and **E)** average staining intensity of positive pixels was quantified in regions around blood vessels (1-9 images per donor) and shown as individual data points. Differences between groups were assessed using a mixed-model analysis. Data are presented as median ± IQR. Never-smoker controls: n = 9, ex-smoker controls: n=9, IPF: n = 12. COL6α1: type VI collagen α1-chain, IPF: idiopathic pulmonary fibrosis, IQR: interquartile range.

Similar to what was observed for COL6α1, when evaluating COL6α2 staining, localization was primarily seen around the airways and blood vessels, with a less distinct distribution in the parenchyma in controls (**Figure 5A-B)**. In IPF, there was strong staining throughout the tissue, including around the airways and blood vessels (**Figure 5C**). When analyzing whole tissue sections of COL6α2 staining, no differences were observed in the proportion of positively stained tissue area (**Figure 5E**), however, the average staining intensity of positive pixels was lower in IPF compared with never-smoker and ex-smoker controls (p=0.0099 and p=0.0066, respectively, **Supplementary Figure 3E**). In the parenchymal regions, there were no differences in the COL6α2 proportion or the average pixel staining intensity (**Supplementary Figure 3B & F**). Around the airways, both the proportion of COL6α2 and the average positive pixel staining intensity were lower in IPF compared with never-smoker (%area: p=0.033, intensity: p=0.028) and ex-smoker controls (%area: p=0.003, intensity: p=0.004) (**Figure 5D** and **Supplementary Figure 3G**). Similarly, in the blood vessels, there was a significantly lower proportion of COL6α2 and average pixel staining intensity in IPF compared with controls (never-smoker: %area: p=0.012, intensity: p=0.028, ex-smoker: %area: p<0.001, intensity: p=0.004, **Figure 5F** and **Supplementary Figure 3H**). The proportion of COL6α2 was also higher in ex-smokers compared with never-smoker controls (p=0.038, **Supplementary Figure 3D**). Taken together, COL6α2 tissue changes were most pronounced in IPF airways and blood vessels compared with controls.

**Figure 5.**
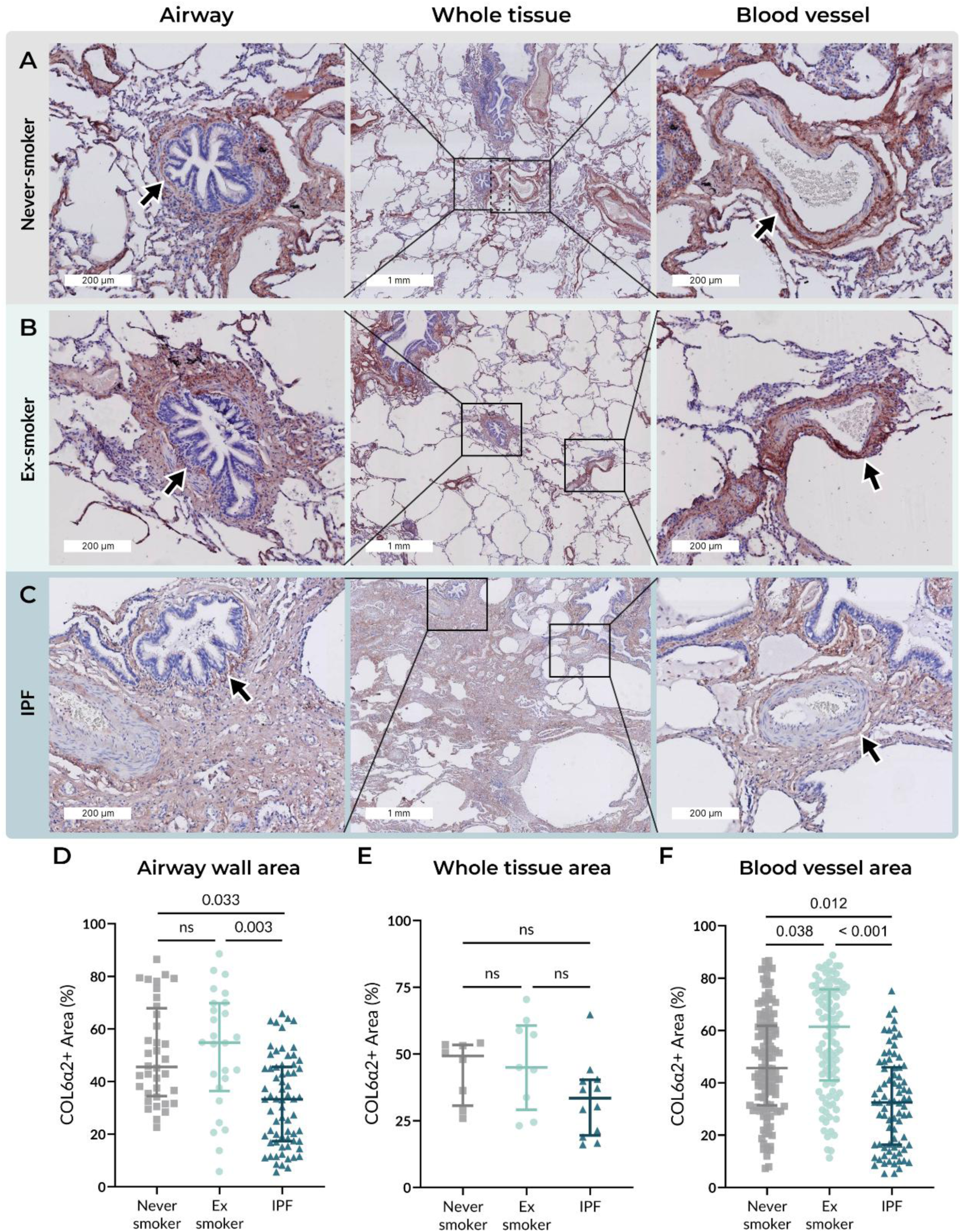
IPF airways and vessels have a lower proportion of COL6α2 compared with controls. Images of sections of lung tissue stained for COL6α2 were digitally quantified. Representative tissue images from **A)** never-smoker control, **B)** ex-smoker control, and **C)** IPF. The images in the center are of whole tissue and have a scale bar of 1 mm. The images to the left zoom in on an airway, and the images to the right zoom in on a blood vessel. The zoomed images have a scale bar of 200 µm. The arrows indicate airways on the left images and blood vessels on the right images. Contrast and saturation were digitally increased by 5% for all images to enhance visibility. **D & F)** Positively-stained area (%) was quantified in airway walls and regions around blood vessels, respectively, (1-9 images per donor) and shown as individual data points and differences between groups were assessed using a mixed-model analysis. Data are presented as median ± IQR. **E)** Each datapoint represents an individual donor. Positively-stained area (%) was quantified in whole tissue, analyzed with one-way ANOVA, and presented as mean ± SD. Never-smoker controls: n = 9, ex-smoker controls: n=9, IPF: n = 12. COL6α2: type VI collagen α2-chain, IPF: idiopathic pulmonary fibrosis, IQR: interquartile range, SD: standard deviation.

### COL6 production is proportionally lower in IPF airways

To better understand COL6 turnover in lung tissue, IHC staining was performed in controls and IPF tissue using antibodies specific for COL6 production (PRO-C6) and degradation (C6M) fragments.

In control lungs, PRO-C6 primarily localized around structural ECM-rich areas, such as airways and blood vessels (**Figure 6A-B**). In IPF tissue, staining in the airways was weak, whereas blood vessels had stronger positive staining (**Figure 6C**). Additionally, heterogeneous staining was observed in fibrotic interstitial regions and PRO-C6 was present in fibroblastic foci (**Supplementary Figure 4**). PRO-C6 image analysis revealed that a lower proportion of the tissue area was occupied by PRO-C6 in IPF airways compared with never-smoker and ex-smoker controls (p=0.0076 and p=0.0002, respectively, **Figure 6D**). No differences were observed in the proportion of tissue occupied by PRO-C6 in other tissue compartments, nor were there any differences in the average positive pixel staining intensities (**Figure 6E** and **Supplementary Figure 5**). In summary, PRO-C6 tissue changes were most pronounced in IPF airways where the tissue had proportionally lower levels compared with controls.

**Figure 6.**
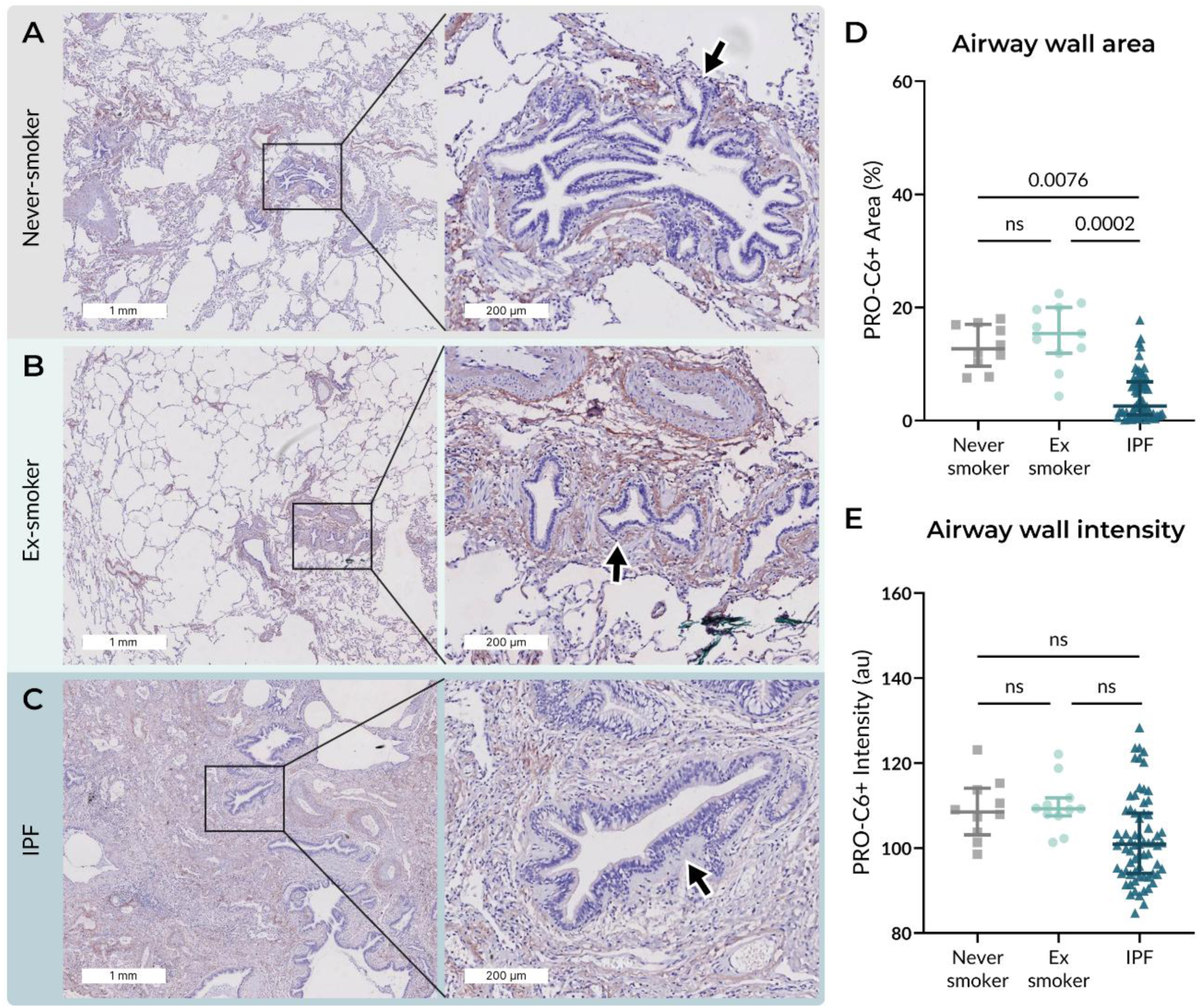
PRO-C6 proportion is lower around IPF airways compared with controls. Images of sections of lung tissue stained for PRO-C6 were digitally quantified. Representative tissue images from **A)** never-smoker control, **B)** ex-smoker control, and **C)** IPF. Scale bars: 1 mm for images on the left; 200 µm for images on the right. The arrows indicate airways. Contrast and saturation were digitally increased by 5% for all images to enhance visibility. **D)** Positively-stained area (%) and **E)** average staining intensity of positive pixels was quantified in airway walls (1-9 images per donor) and shown as individual data points and differences between groups were assessed using a mixed-model analysis. Data are presented as median ± IQR. Never-smoker controls: n = 3, ex-smoker controls: n=5, IPF: n = 10. PRO-C6: type VI collagen production, IPF: idiopathic pulmonary fibrosis, IQR: interquartile range.

In contrast, C6M was widely present throughout the tissue regions in all groups, and it appeared to localize within airway epithelium (**Figure 7A-C**). Analysis of C6M revealed proportionally lower positively stained tissue in IPF compared with never-smoker controls across all tissue compartments (whole tissue: p=0.0008, **Figure 7D**, parenchyma: p=0.0006, airway wall: p=0.0032, blood vessels: p=0.0104, **Supplementary Figure 6B-D**). Similar differences were observed for the average positive pixel staining intensity (whole tissue: p=0.0015, **Figure 7E**, parenchyma: p=0.0059, airway wall: p=0.0161, blood vessels: p=0.0079, **Supplementary Figure 6E-H**). No differences in the proportion of C6M or average pixel staining intensity were found between ex-smoker controls and IPF tissue (**Supplementary Figure 6**). Notably, a significant lower proportion of C6M and average pixel staining intensity were found in the whole tissue, parenchyma, and blood vessels of ex-smoker controls compared with never-smoker controls (whole tissue: area: p=0.0165, intensity: p=0.0389, parenchyma: area: p=0.0199, intensity: p=0.0402, blood vessels: area: p=0.0450, intensity: p=0.0171, **Supplementary Figure 6A-B, D-F & H**). Overall, both the protein proportion and average positive pixel staining intensity of C6M were lower in all IPF tissue compartments compared with never-smoker but not ex-smoker controls.

**Figure 7.**
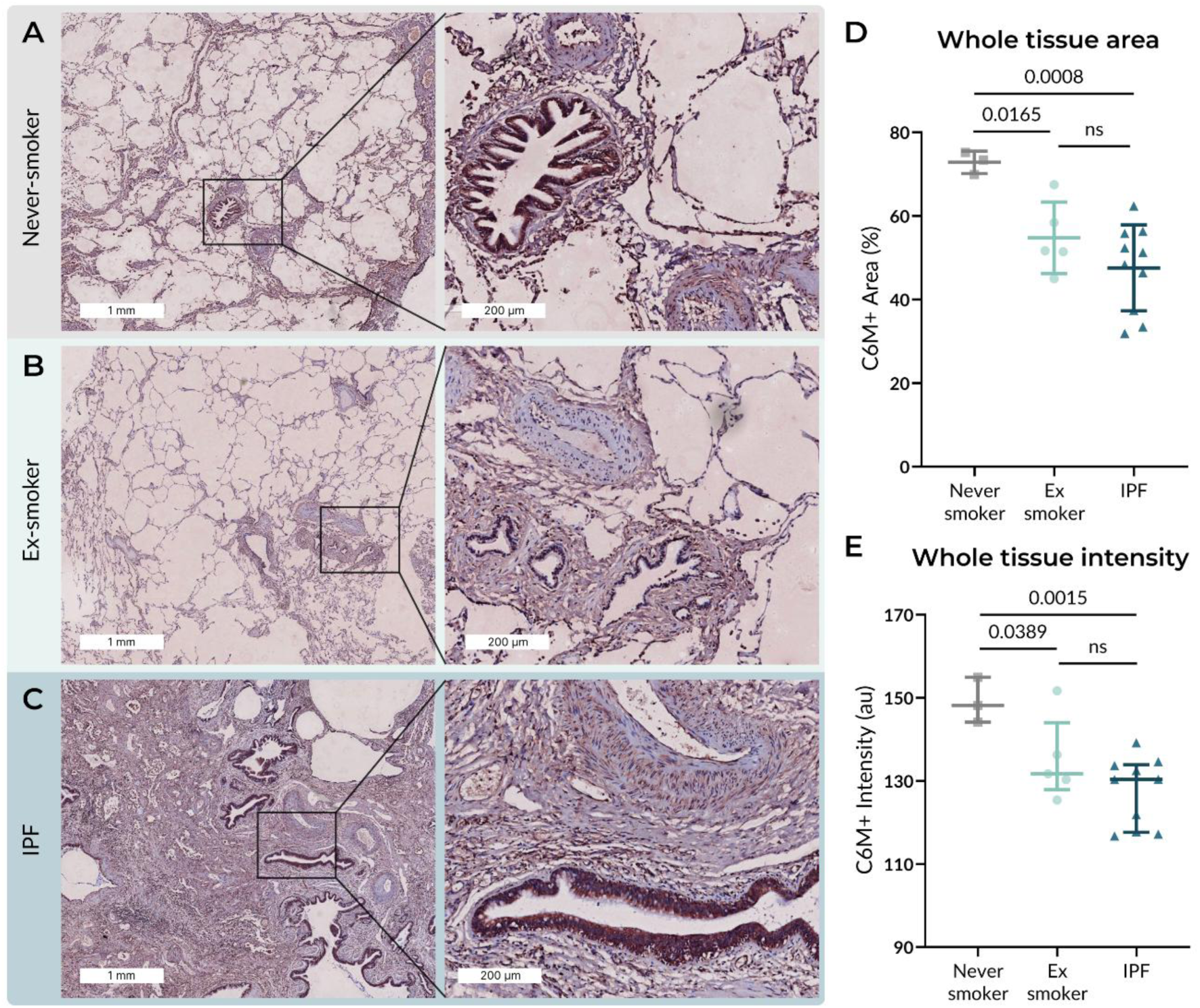
C6M is proportionally lower in IPF lung tissue compared with controls. Digitalized images of sections of lung tissue stained for C6M were digitally quantified. Representative tissue images from **A)** never-smoker control, **B)** ex-smoker control, and **C)** IPF. Scale bars: 1 mm for images on the left; 200 µm for images on the right. Contrast and saturation were digitally increased by 5% for all images to enhance visibility. **D)** Positively-stained area (%) and **E)** average staining intensity of positive pixels was quantified in whole tissue and analyzed with one-way ANOVA. Data are presented as mean ± SD. Never-smoker controls: n = 3, ex-smoker controls: n=5, IPF: n = 10. C6M: type VI collagen degradation, IPF: idiopathic pulmonary fibrosis, SD: standard deviation.

In summary, quantitative analysis of IHC stained lung tissue revealed that C6M was proportionally lower in IPF compared with never-smoker controls in both whole tissue and different lung tissue compartments (**Figure 8**). As for COL6α1, COL6α2, and PRO-C6, no differences in protein or peptide proportion were found between never-smoker controls and IPF in whole tissue and parenchyma. When investigating the airway wall, COL6α2 and PRO-C6 proportions were significantly lower in IPF compared with never-smoker and ex-smoker controls. Around the blood vessels, COL6α1 and COL6α2 were proportionally lower in IPF compared with never-smoker and ex-smoker controls. COL6α1 had a lower average positive pixel staining intensity in IPF compared with never-smoker and ex-smoker controls in both whole tissue, parenchyma, and blood vessels. COL6α2 also had a lower positive pixel staining intensity in IPF compared with both never-smoker and ex-smoker controls in the airway wall and blood vessel regions, and in the whole tissue compared with ex-smoker controls. PRO-C6 did not differ between groups in any lung tissue compartments, whereas C6M had a lower average staining intensity in IPF compared with never-smoker controls across all lung tissue compartments.

**Figure 8.**
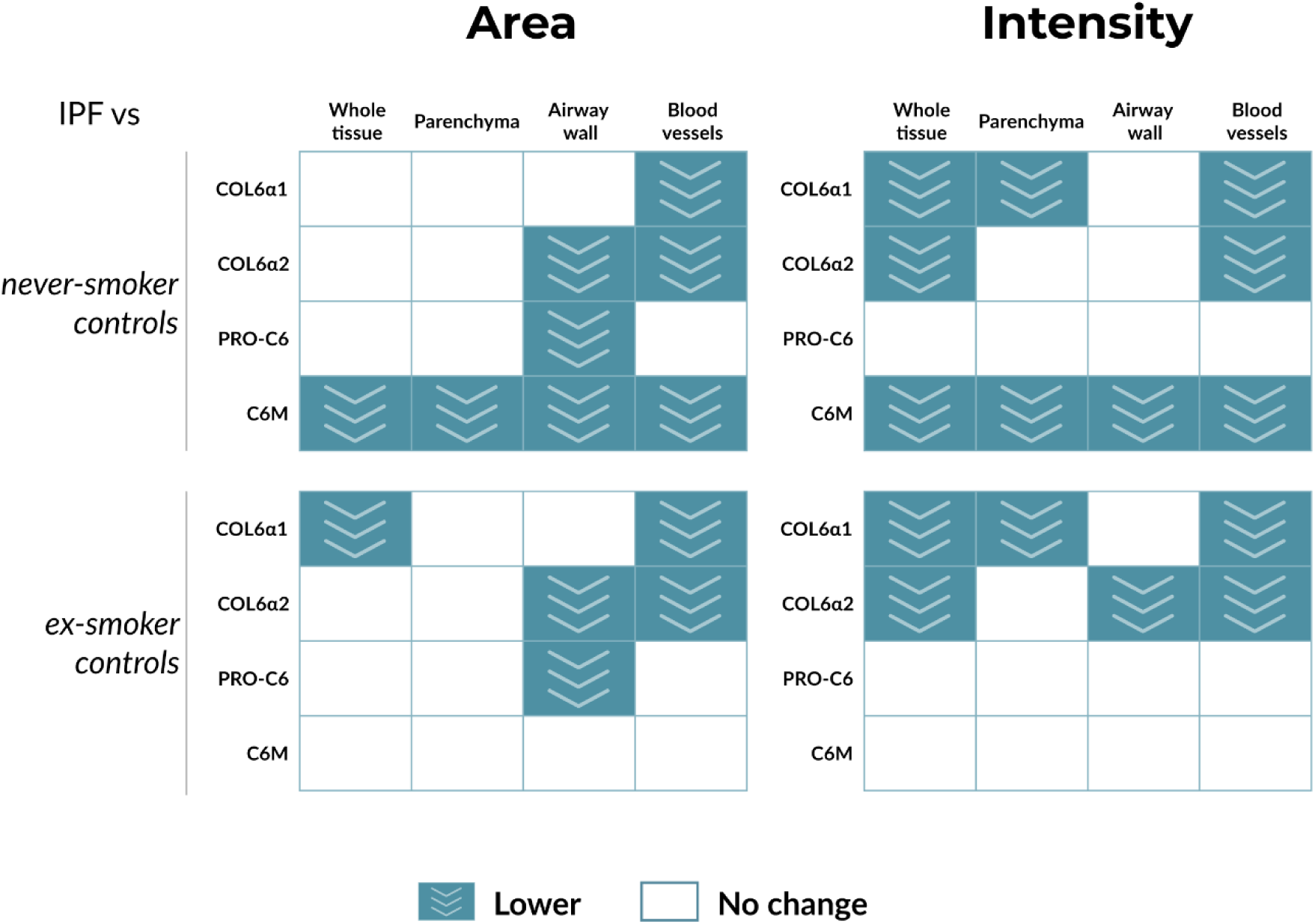
Quantification of COL6 staining in IPF and controls. The proportion of positively-stained tissue area (the left panels) and the average positive pixel staining intensity (the right panels) are shown as a comparison between IPF and never-smoker controls (the top panels) and ex-smoker controls (the bottom panels). Within each panel, proportions of COL6α1, COL6α2, PRO-C6, and C6M are reflected for each tissue compartment: whole tissue, parenchyma, airway wall, and blood vessels. The filled rectangles with arrows pointing downwards represent lower proportions, whereas empty rectangles with no arrows represent no change.

### COL6 and PRO-C6 promote viability in lung fibroblasts

Finally, we wanted to characterize the effects of COL6 and fragments released during its production (PRO-C6) and degradation (C6M) on the viability and apoptosis of non-diseased control lung fibroblasts, bronchial epithelial cells, and pulmonary microvascular endothelial cells. These data showed that the cell viability was higher in the fibroblasts treated with COL6 compared with untreated fibroblasts (p=0.002, **Figure 9A**). Similarly, treatment with PRO-C6 resulted in higher fibroblasts viability compared with untreated controls (p=0.0021, **Figure 9B**). In contrast, C6M treatment did not result in detectable changes in the fibroblast viability (**Figure 9C**). Similarly to the effects on fibroblasts, both epithelial and endothelial cells showed increased viability trends when treated with COL6 and PRO-C6, as well as with C6M (**Supplementary Figure 7A-C** and **8A-C**). When assessing the potential of COL6 and its fragments in inducing apoptosis through caspase activity, none of the treatments resulted in a detectable change in fibroblasts (**Figure 9D-F**). Similarly, no effects were seen in epithelial cells (**Supplementary Figure 7D-F**), whereas endothelial cells showed a trend towards decreased apoptotic activity in the presence of COL6 and PRO-C6 (**Supplementary Figure 8D-F**).

**Figure 9.**
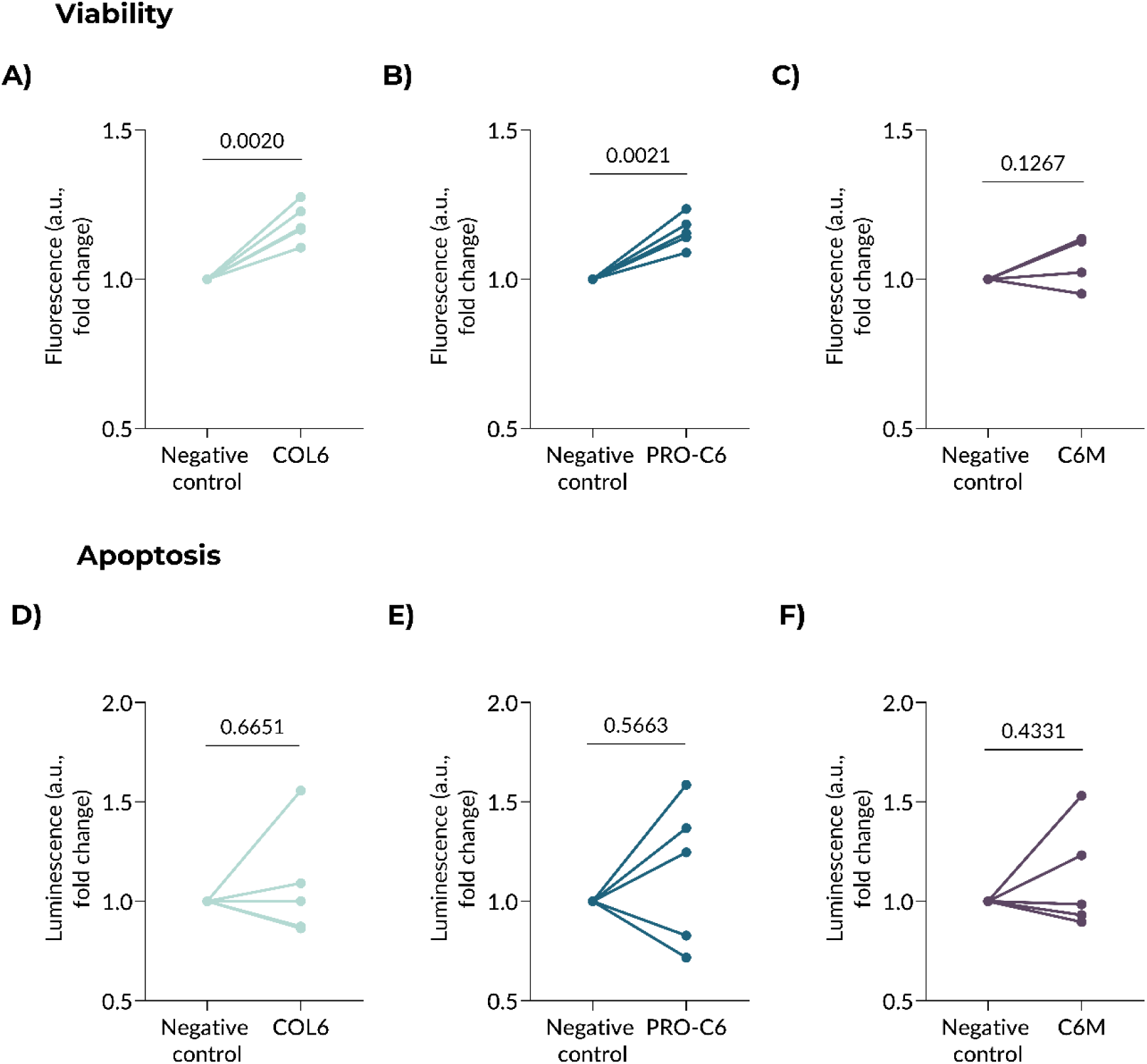
COL6 and PRO-C6 increased fibroblast viability. Primary human lung fibroblasts were cultured without (negative control) or with 8 ng/μL of COL6, PRO-C6, or C6M for 24h and cell viability and apoptosis were measured using ApoLive-Glo Multiplex Assay. The number of viable cells (viability) was represented as the fold change in fluorescence signal compared with untreated negative control for the groups: **A)** COL6, **B)** PRO-C6, and **C)** C6M. Caspase activity 3/7 (apoptosis) was represented as the fold change in luminescence compared with the negative control for the groups: **D)** COL6, **E)** PRO-C6, and **F)** C6M. Each datapoint represents one biological donor (n=5). Log-transformed data were analyzed with a paired t-test and presented as fold change from the negative control. a.u.: arbitrary unit, C6M: type VI collagen degradation, COL6: type VI collagen, PRO-C6: type VI collagen production.

## Discussion

In this study, we described how COL6 gene and protein expression are altered in IPF, including the turnover and spatial distribution of specific COL6 remodeling fragments. Genes associated with the five functional COL6 α chains had higher expression in IPF lungs compared with controls; while the protein proportion of COL6α1, COL6α2, and COL6 production fragment PRO-C6 in the airways and/or blood vessels were lower in IPF lungs compared with never-smoker control lungs. Consistently lower proportions of COL6 degradation fragment C6M across different lung tissue compartments indicate that COL6 degradation occurs in the whole lung. When tested *in vitro* on primary lung cells, the viability of fibroblasts was promoted by COL6 and its production fragment PRO-C6. Taken together, our results illustrate how COL6 and its production fragment are proportionally lower in specific IPF lung tissue compartments, primarily around the airways and blood vessels, which suggests a potential association with the BM. To the best of our knowledge, this report is the first to characterize COL6 and its remodeling fragments in IPF lung tissue.

COL6 is one of the most common lung ECM proteins, and it has recently been shown to remain consistently abundant in lung tissue derived from control donors and patients with lung fibrosis or chronic obstructive pulmonary disease [8]. COL6 protein primarily localizes in blood vessels, bronchi, bronchioles, and the alveolar interstitium [13]. In the case of fibrosis, COL6 protein also localizes within the myofibroblast core of fibroblastic foci [46]. These earlier findings are also corroborated at the RNA level: our transcriptomics analysis of publicly available datasets revealed that in fibrotic lung samples, all five functional genes encoding the COL6 α chains showed higher expression in mesenchymal cells. Mesenchymal cells are major ECM producers, and these data are consistent with previous reports showing increased COL6 gene expression in IPF lungs compared with non-fibrotic lungs [47]. Moreover, our analysis of epithelial, endothelial, and immune cells showed higher expression of the genes encoding the COL6 α1, α2, and α3 chains. It remains unclear whether these higher gene expression levels reflect a response to the fibrotic microenvironment or actively contribute to alterations of the microenvironment.

COL6 protein is found at various locations throughout the lung tissue. Our analysis of the IHC images revealed that both COL6α1 and COL6α2 were present in parenchyma, airway walls, and blood vessels in non-diseased control lung tissue. This aligns with previous findings by Specks *et al.* who showed that COL6 localizes in the interstitial space, bronchial walls, and vasculature of control lungs [13]. Furthermore, when we quantified specific structural compartments, the proportion of COL6α2 was lower in IPF airways compared with controls, and the levels of both COL6α1 and COL6α2 were proportionally lower in IPF blood vessels. A common feature in both airways and blood vessels is the BM, which is essential for gas transfer. This shift in COL6 spatial distribution in fibrotic tissue thus suggests a change in the ECM composition around epithelial and endothelial cells, possibly disrupting the link between the BM and IM. Importantly, such a disruption could impair the epithelial/endothelial-fibroblast crosstalk, which plays a key role in epithelial and endothelial regeneration and during the fibrotic response [48–50].

In the fibrotic lung, another key feature is increased crosslinks and aberrant ECM remodeling, which together with excessive deposition of ECM, increases the lung stiffness [7]. In cancer, which is commonly associated with fibrosis around tumor cells, glioblastoma cell cultures secrete higher levels of COL6, correlating with increased ECM stiffness and tumor invasion [51]. The localization of COL6 in the interstitium of dense fibrotic tissue raises the question of whether COL6 contributes to increased ECM stiffness, or whether increased ECM stiffness induces COL6 synthesis. Nevertheless, increased ECM stiffness can drive a pro-fibrotic fibroblast phenotype characterized by apoptosis resistance, high rates of proliferation, and ECM production [7,52]. Additionally, COL6 contains a large amount of von Willebrand factor A-like domains, which can bind to platelets and activate them [53]. Their subsequent release of transforming growth factor beta (TGF-β) and platelet-derived growth factor (PDGF) can further promote fibroblast activation and ECM production, in a self-perpetuating loop, making COL6 not just a product of fibrosis, but also a driver of it, in a similar fashion that tissue stiffness drives further fibrotic responses.

Emerging evidence points at how ECM organization and topography play a role in IPF progression [3,54]. Macrophages encountering fibrous collagen fibers had a faster migratory rate than those on globular collagen fibers [55]. While the topological arrangement of the ECM was not explored in the current study, it can be speculated that an altered COL6 localization and proportion within the tissue would result in an abnormal structural organization of the ECM, which then could influence infiltrating immune cells. COL6 has been shown to suppress fibronectin induced cellular migration in the developing gut [56]. Thus, proportionally lower levels of COL6 at the BM-IM interface could potentially facilitate cell migration into the tissue, particularly where this change is around the blood vessels. Cell-cell interactions can also affect immune cell migration [2]. Thus, if reduced COL6 or altered spatial distribution influences epithelial-fibroblast or endothelial-fibroblast crosstalk, this might in turn affect the migration of immune cells.

Previous studies have shown that patients with IPF have extensive vascular remodeling, with thickened vascular walls and an altered ECM deposition [57,58]. Thus, it may be speculated whether the altered ECM deposition in blood vessels is characterized by lower levels of COL6 as found in the current study. In the skeletal muscle of mouse, COL6α3 colocalized with an endothelial marker, suggesting that COL6, at least in part, associated with blood vessels [59]. In agreement with our findings, a recent study showed that upon exposure to hypoxia, there was decrease in COL6 protein in the tunica media of blood vessels of rat lungs [60]. In another study, COL15A1 positive vessels could replace other lung-specific vasculatures upon regeneration after injury [61]. Accordingly, it is plausible that the regeneration of the vasculature or the failure to regenerate is characterized by an altered ECM composition, including lower levels of COL6, perhaps by being replaced by other collagens.

Production of COL6 (PRO-C6) was localized in the same structural ECM-rich areas containing BM as COL6α1 and α2, namely around airways and blood vessels, and had a proportionally lower presence in IPF airways. This lower relative amount could suggest that COL6 production in the fibrotic airways is disturbed. Additionally, the PRO-C6 production fragment was present in fibroblastic foci, indicating active COL6 production by fibroblasts in IPF. PRO-C6 has also been detected in fibrotic tissue in other organs, such as the kidney and liver [62,63], supporting its relevance beyond pulmonary fibrosis. Importantly, the PRO-C6 antibody targets a part of the COL6α3 C5 domain which is cleaved of during COL6 maturation, namely the signaling fragment endotrophin. Endotrophin has pro-fibrotic properties and is known to be associated with IPF progression as well as increased expression of ECM and ECM regulating genes [18–20,27–29]. Thus, it is likely that COL6 contributes to the fibrotic response through the release of endotrophin, and that this specific COL6 fragment (PRO-C6) has the potential to induce cellular responses, different than those of intact COL6. A recent study confirmed that the C-terminal part of COL6α3, including endotrophin, was cleaved off after microfibril formation prior to incorporation into the tissue [59]. Similar to our current study, this study also identified the fragment in tissue staining and concluded that its release into circulation was not due to increased proteolysis.

COL6 degradation is part of regular tissue homeostasis, which is mainly performed by the gelatinases MMP-2 and MMP-9 [64]. Contrary to the more localized COL6 production fragment (PRO-C6), the MMP-mediated COL6 degradation fragment (C6M) showed strong and widespread staining throughout the tissue, indicating that COL6 degradation occurs in the whole lung. C6M is released from inflamed and fibrotic tissue and has previously been associated with chronic inflammation [65]. Interestingly, we found lower proportional levels of tissue-associated C6M around the blood vessels of ex-smoker controls compared with never-smoker controls, while there were no differences between ex-smoker and IPF tissues proportions. This difference was not observed for COL6α1, the parent molecule from which the C6M fragment originates from. While previous studies show contradictory results on the effect of smoking cessation, most suggest that airway inflammatory and structural vascular changes induced by smoking persist [66–68]. These observations raise an interesting question as to whether the vascular changes that occur in smokers’ lungs influence the tissue collagen composition or availability of the antibody-binding domain of COL6, making it more difficult to degrade. Taken together, these findings indicate region-specific alterations in COL6 turnover in IPF, with ongoing COL6 remodeling in the IM, and airways showing the most pronounced changes in remodeling.

The importance of COL6 in regulating biological mechanisms has been demonstrated in several previous studies. COL6 has versatile roles in different tissues such as the skin, nervous system, infarcted heart, and lung [9]. In our study, COL6 and its production fragment PRO-C6 (endotrophin), but not degradation fragment (C6M), increased the viability of primary lung fibroblasts. Similar trends were observed in primary bronchial epithelial and pulmonary micro endothelial cells, with analyses remaining descriptive due to the limited number of donors. These findings align with previous reports suggesting that endotrophin can amplify fibrotic processes, potentially through direct effects on fibroblasts [27,69]. While these results highlight that both COL6 and its production fragment can promote cell viability, it remains unclear how their change in protein/peptide proportion within the tissue influence fibroblast viability. We also investigated the influence of COL6 and its fragments on apoptosis, as several studies have demonstrated a role for COL6 in the regulation of apoptotic processes. Knockout of *Col6a1* in mouse embryonic fibroblasts led to spontaneous apoptosis, which could be rescued when the cells were cultured on COL6-coated, but not COL1-coated, plates [25]. It has also been shown that COL6 prevents ultraviolet-induced apoptosis in primary mouse hippocampal neurons [70]. In contrast, myocyte apoptosis was alleviated in *Col6a1*-knockout mice following myocardial infarction [71]. In our study, treatment with COL6, PRO-C6, or C6M did not affect caspase activity in primary lung fibroblasts, and a similar trend was observed in epithelial cells. In contrast, apoptotic activity in endothelial cells showed a decreasing trend; however, this observation would require further validation in a larger sample set.

Our findings have potential implications for future studies and for how we approach IPF and its progression, although several limitations should be acknowledged. The cell culture experiments presented here were exploratory and conducted in 2D systems. Consequently, future studies using more sophisticated and biologically relevant models [72] will be important in elucidating the influence of COL6 and its remodeling fragments. Another limitation of our study is the lack of information on the medication history of the donors. We do not know whether anti-fibrotic therapies were used, therefore it is not possible to know whether heterogeneity in the staining data may reflect patients’ medication history. Further analysis of the history of fibrotic therapies related to COL6 and its fragments that are anchored in the tissue should be a focus for future research. In addition, measuring COL6 fragments in blood samples matched with lung tissue staining from the same patients, would provide a more comprehensive understanding of COL6 dynamics in IPF, which could ultimately help in developing more targeted methods for disease monitoring and treatment.

In conclusion, our study highlights the dynamics of COL6 remodeling in the lung ECM and its dysregulation in IPF. We demonstrated that COL6 α1 and α2 chains are found in proportionally lower amounts around airways and blood vessels of IPF lungs compared with non-fibrotic lungs. As these structures are closely associated with the BM, our results on altered COL6 remodeling suggest potential implications on BM integrity, cell-cell interactions, and immune cell infiltration. Furthermore, the distinct localizations of fragments associated with COL6 production (PRO-C6) and degradation (C6M) illustrate the complexity of COL6 turnover in fibrotic tissue. Together with the functional effects of COL6 and PRO-C6 through increased primary lung fibroblast viability, our results show how COL6 presence and its altered turnover in lung tissue influence fibroblast survival and in turn contribute to continued fibrotic processes. Collectively, our characterizations provide novel insights into the spatial and functional involvement of COL6 in IPF.

## Acknowledgements

The stainings of COL6α1 and COL6α2 on lung tissue analyzed in this manuscript were conducted as part of the HOLLAND (HistopathOLogy of Lung Aging aNd COPD) project. The HOLLAND project was initiated and supervised by Corry-Anke Brandsma, Wim Timens, and Janette Burgess. Technical support was provided by Marjan Reinders-Luinge, Anja Bakker, and Theo Borghuis, and image analyses pipelines were developed by Theo Borghuis, Maunick Lefin Koloko Ngassie, and Niek Bekker. Authors thank Mr. Albano Tosato for the assistance in figure preparation and Dr Meike Zwager for reviewing the PRO-C6 staining in fibrotic foci in IPF lung tissue sections.

## Abbreviations

AEGM: Airway Epithelial Growth Medium
BM: Basement Membrane
C6M: Type VI Collagen Degradation
COL6: Type VI Collagen
DMEM: Dulbecco’s Modified Eagle Medium
ECM: Extracellular Matrix
FBS: Fetal Bovine Serum
FEV1: Forced Expiratory Volume in 1 Second
IHC: Immunohistochemistry
IM: Interstitial Matrix
IPF: Idiopathic Pulmonary Fibrosis
P/S: Penicillin–Streptomycin
PRO-C6: Type VI Collagen Production
scRNAseq: Single-Cell RNA Sequencing
UMCG: University Medical Center Groningen.

## Data availability

Data are available upon reasonable request.

## Funding

This study has been funded by the Danish Research Foundation and Noordelijke Cara Stichting, Netherlands.

## Disclosures

H.W. Breisnes, M. Karsdal, D.J. Leeming, F.B. Simões, and J.M.B. Sand are employees and may be shareholders of Nordic Bioscience, a company involved in the discovery and development of biomarkers. M. Nizamoglu and J. K. Burgess received unrestricted research funds from Boehringer Ingelheim.

## Author contributions

Conceptualization: HWB, MN, MLKN, JMBS, JKB. Methodology: HWB, MN, MLKN, TB, CAB, JKB. Formal analysis: HWB, MN, MLKN, MRJ, RM. Investigation: HWB, MN, MLKN, TB, MRJ, RM. Resources: TK. Writing – Original draft: HWB, MN, MLKN, MRJ. Writing – Review & Editing: HWB, MN, MLKN, TB, MRJ, RM, TK, CAB, SFT, MK, DJL, FBS, JMBS, JKB. Visualization: HWB, MN. Supervision: JMBS, JKB.

## Notes

### Competing Interest Statement

The authors have declared no competing interest.

