## Supplementary Material for "Type VI collagen is proportionally lower around airways and blood vessels in idiopathic pulmonary fibrosis"

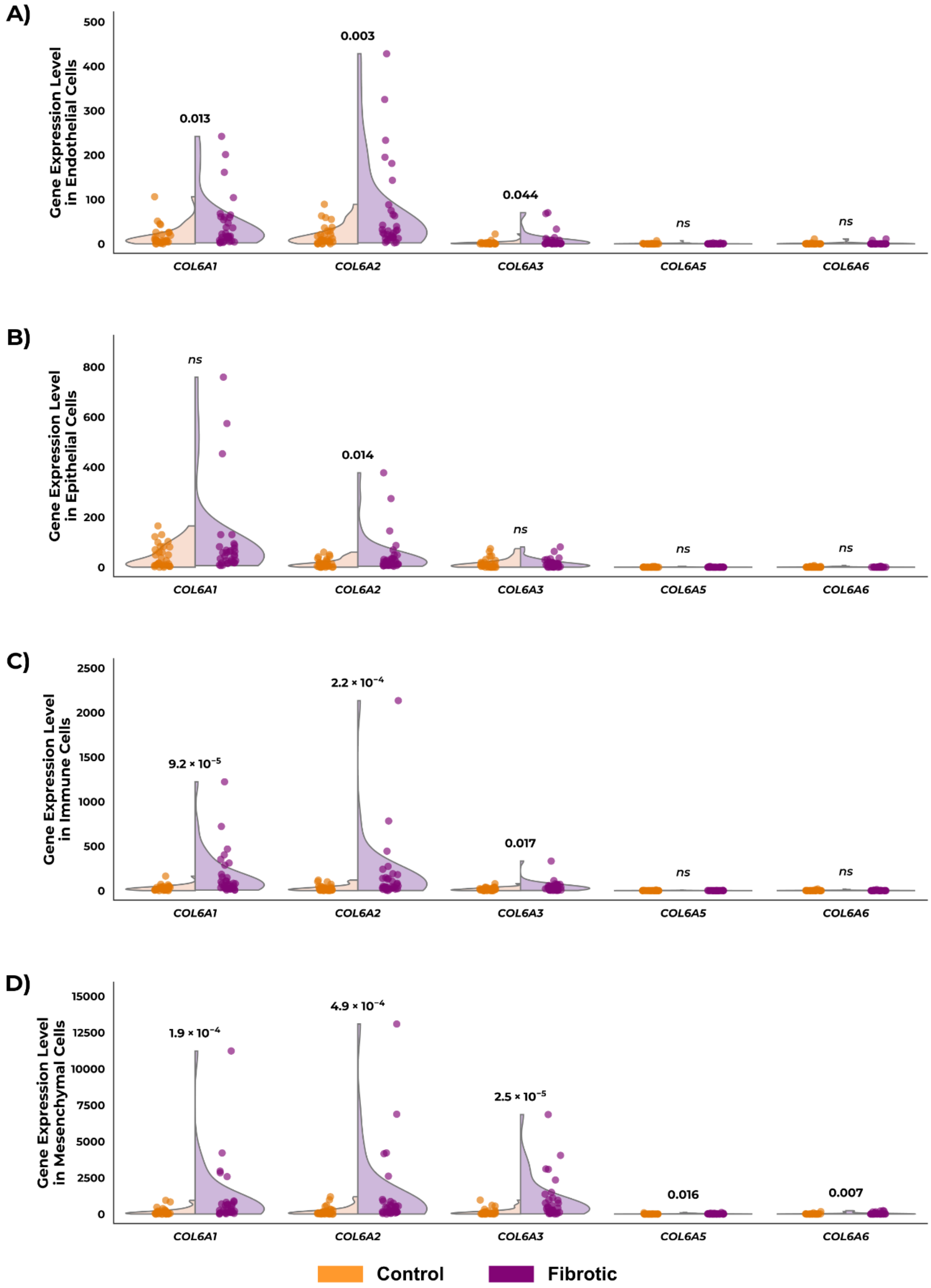

***Supplementary Figure 1.* COL6A1-A3, A5*, and* A6 *are overexpressed in fibrotic lung cells compared with controls***. *Pseudobulk analysis of gene expression profiles for* COL6A1-A3, A5 *and* A6 *was performed using publicly available single cell RNA-sequencing data [34] to compare expression in* ***A)*** *endothelial cells,* ***B)*** *epithelial cells,* ***C)*** *immune cells, and* ***D)*** *mesenchymal cells. n=30 controls, n=32 fibrotic donors. Each dot represents one biological donor. Statistical comparisons between control and fibrotic donors per COL6 encoding gene were performed using Mann-Whitney test. COL6: type VI collagen.*

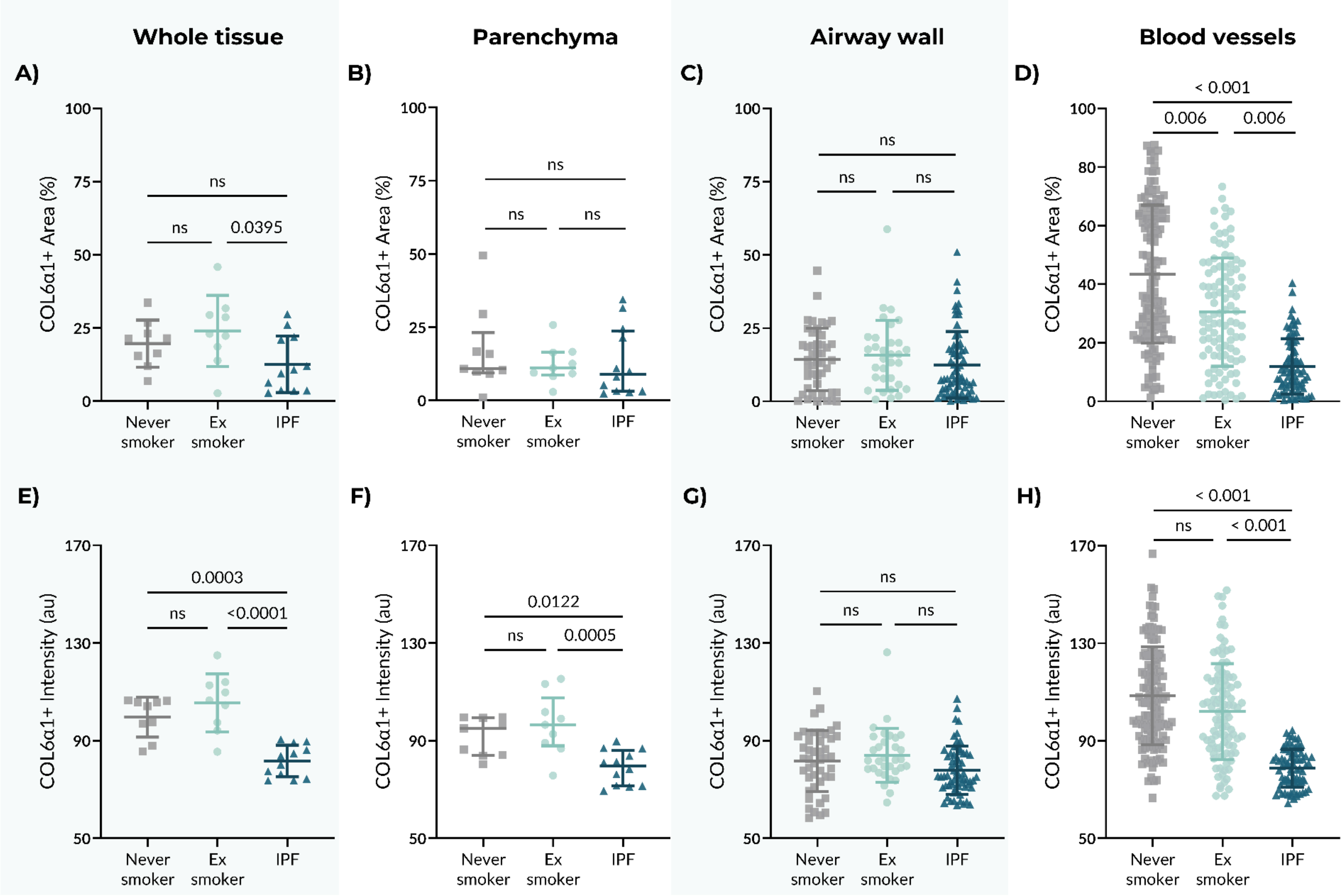

Whole tissue
Parenchyma
Airway wall
Blood vessels
A)
B)
C)
D)
E)
F)
G)
H)
COL6α1+ Area (%)
COL6α1+ Intensity (au)
Never smoker
Ex smoker
IPF
ns
0.0395
0.0003
<0.0001
0.0122
0.0005
< 0.001
0.006


9

***Supplementary Figure 2. Image analysis on immunohistochemical staining for COL6α1 in human lung tissue from controls and patients with IPF****. The area that was positively-stained (%) for COL6a1 was quantified in* ***A****) whole tissue,* ***B****) parenchyma,* ***C****) airway wall, and* ***D)*** *blood vessels. The average staining intensity of pixel positive for COL6a1 (arbitrary units [au]) was quantified in* ***E****) whole tissue,* ***F****) parenchyma,* ***G****) airway wall, and* ***H)*** *blood vessels. For panels* ***A-B*** *and* ***E-F****, each datapoint represents an individual donor; panels* ***A*** *and* ***E-F*** *were analyzed using one-way ANOVA and shown as mean ± SD, whereas panel* ***B*** *was analyzed using the Kruskal-Wallis test and shown as median ± IQR. In panels* ***C-D*** *and* ***G-H****, individual airway or blood vessel images (1-9 images per patient) are shown as individual datapoints with median ± IQR, and differences between groups were assessed using a mixed-model analysis to account for multiple images airways per patient. Samples sizes: n = 9 for never-smoker and ex-smoker controls, n = 12 for IPF. COL6α1: type VI collagen α1 chain, IPF: idiopathic pulmonary fibrosis, IQR: interquartile range, SD: standard deviation.*

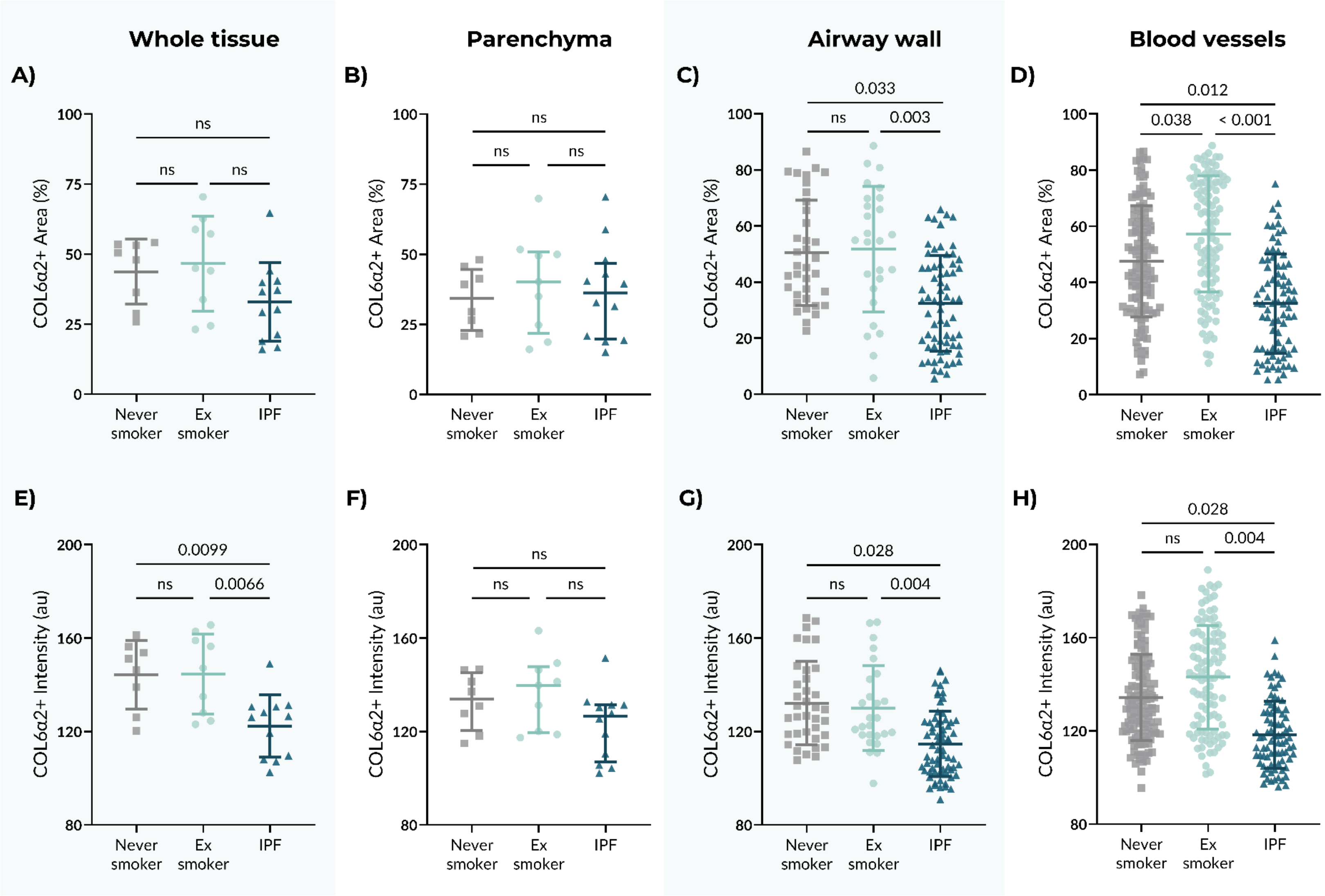
Whole tissue
Parenchyma
Airway wall
Blood vessels
A)
B)
C)
D)
E)
F)
G)
H)
COL6α2+ Area (%)
COL6α2+ Intensity (au)
Never smoker
Ex smoker
IPF
ns
0.033
0.003
0.012
0.038
< 0.001
0.0099
0.0066
0.028
0.004

22

***Supplementary Figure 3. Image analysis on immunohistochemical staining for COL6α2 in human lung tissue from control and patients with IPF.*** *The area that was positively-stained (%) for COL6α2 was quantified in **A**) whole tissue, **B**) parenchyma, **C**) airway wall, and **D)** blood vessels. The average staining intensity of pixel positive for COL6a2 (arbitrary unit [au]) was quantified in **E**) whole tissue, **F)** parenchyma, **G**) airway wall, and **H)** blood vessels. For panels **A-B** and **E-F**, each datapoint represents an individual donor; panels **A** and **E-F** were analyzed using one-way ANOVA and shown as mean ± SD, whereas panel **B** was analyzed using the Kruskal-Wallis test and shown as median ± IQR. In panels **C-D** and **G-H**, individual airway or blood vessel images (1-9 images per patient) are shown as individual datapoints with median ± IQR, and differences between groups were assessed using a mixed-model analysis to account for multiple images airways per patient. Samples sizes: n = 8 for never-smoker controls, n=9 for ex-smoker controls, n = 12 for IPF. COL6α2: type VI collagen α2, IPF: idiopathic pulmonary fibrosis, IQR: interquartile range, SD: standard deviation.*

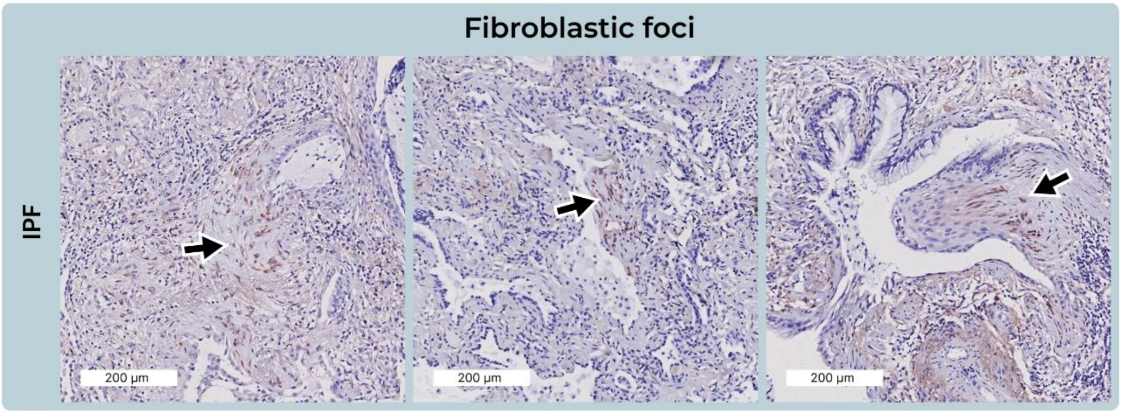


***Supplementary Figure 4. Immunohistochemical staining for PRO-C6 in fibroblastic foci of IPF lung tissue.*** *Representative images of fibroblastic foci in three different IPF tissue donors. Arrows indicate fibroblastic foci. Scale bars: 200 µm. Red/brown color is NovaRed detection of PRO-C6, whereas blue color is hematoxylin counter staining. Contrast and saturation were digitally increased by 5% for all images to enhance visibility.*

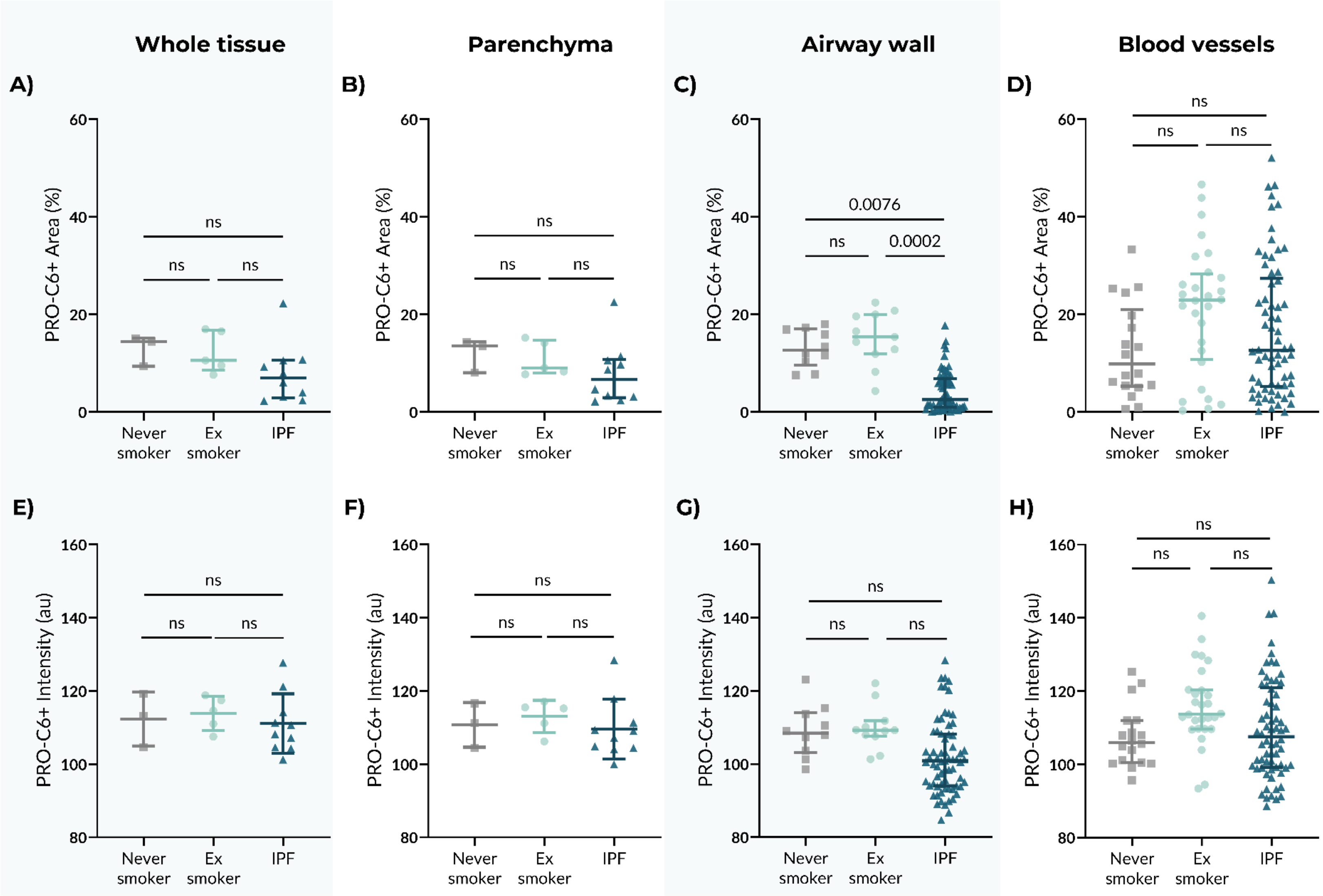
Whole tissue
Parenchyma
Airway wall
Blood vessels
A)
B)
C)
D)
E)
F)
G)
H)
PRO-C6+ Area (%)
PRO-C6+ Intensity (au)
Never smoker
Ex smoker
IPF
ns
0.0076
0.0002

41

***Supplementary Figure 5. Image analysis on immunohistochemical staining for PRO-C6 in human lung tissue from never-smoker controls, ex-smoker controls, and IPF samples.*** *The area that was positively-stained (%) for PRO-C6 was quantified in* ***A)*** *whole tissue,* ***B)*** *parenchyma,* ***C)*** *airway wall, and* ***D)*** *blood vessels. The average staining intensity of pixel positive for PRO-C6 (arbitrary unit [au]) was quantified in* ***E)*** *whole tissue,* ***F)*** *parenchyma,* ***G)*** *airway wall, and* ***H)*** *blood vessels. For panels* ***A-B*** *and* ***E-F****, each datapoint represents an individual donor; data in panels* ***A-B*** *are analyzed with Kruskal-Wallis test and shown as median ± IQR, whereas data in panels* ***E-F*** *are analyzed with one-way ANOVA and shown as mean ± SD. In panels* ***C-D*** *and* ***G-H****, individual airway or blood vessel images (1-9 images per patient) are shown as individual datapoints with median ± IQR, and differences between groups were assessed using a mixed-model analysis to account for multiple features per patient. Never-smoker controls: n = 3, ex-smoker controls: n=5, IPF: n = 10. PRO-C6: type VI collagen production, IPF: idiopathic pulmonary fibrosis, IQR: interquartile range, SD: standard deviation.*

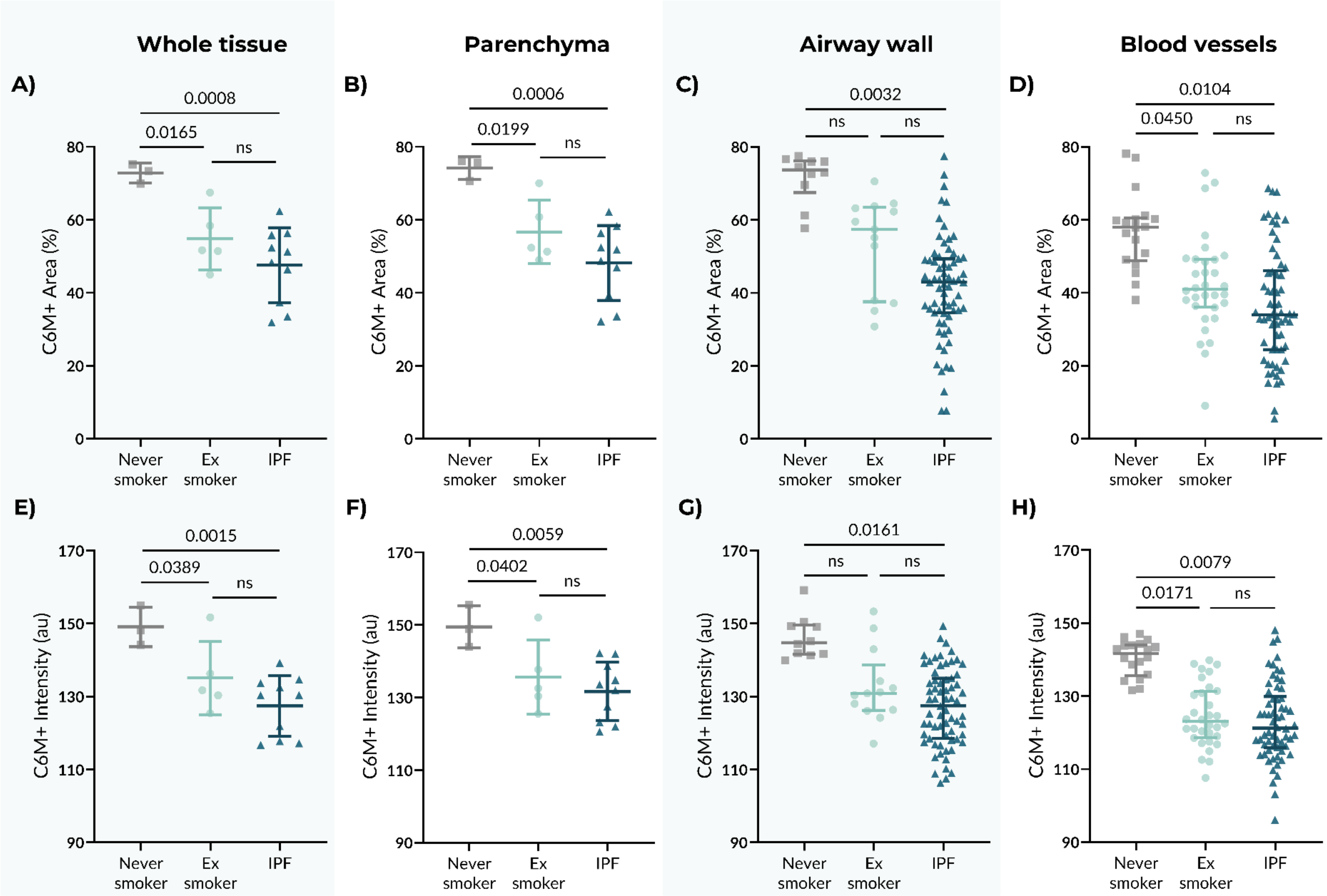
Whole tissue
Parenchyma
Airway wall
Blood vessels
A)
0.0008
0.0165
ns
C6M+ Area (%)
Never smoker
Ex smoker
IPF
B)
0.0006
0.0199
ns
C)
0.0032
ns
ns
D)
0.0104
0.0450
ns
E)
0.0015
0.0389
ns
C6M+ Intensity (au)
F)
0.0059
0.0402
ns
G)
0.0161
ns
ns
H)
0.0079
0.0171
ns

54

***Supplementary Figure 6. Image analysis on immunohistochemical staining for C6M in human lung tissue from never-smoker controls, ex-smoker controls, and IPF samples.*** *The area that was positively-stained (%) for C6M was quantified in **A)** whole tissue, **B)** parenchyma, **C)** airway wall, and **D)** blood vessels. The average staining intensity of pixel positive for C6M (arbitrary unit [au]) was quantified in **E)** whole tissue, **F)** parenchyma, **G)** airway wall, and **H)** blood vessels. For panels **A-B** and **E-F**, each datapoint represents an individual donor, were analyzed with one-way ANOVA, and shown as mean ± SD. In panels **C-D** and **G-H**, individual airway or blood vessel images (1-9 images per patient) are shown as individual datapoints with median ± IQR, and differences between groups were assessed using a mixed-model analysis to account for multiple features per patient. Never-smoker controls: n = 3, ex-smoker controls: n=5, IPF: n = 10. C6M: MMP-mediated type VI collagen degradation, IPF: idiopathic pulmonary fibrosis, IQR: interquartile range, SD: standard deviation.*

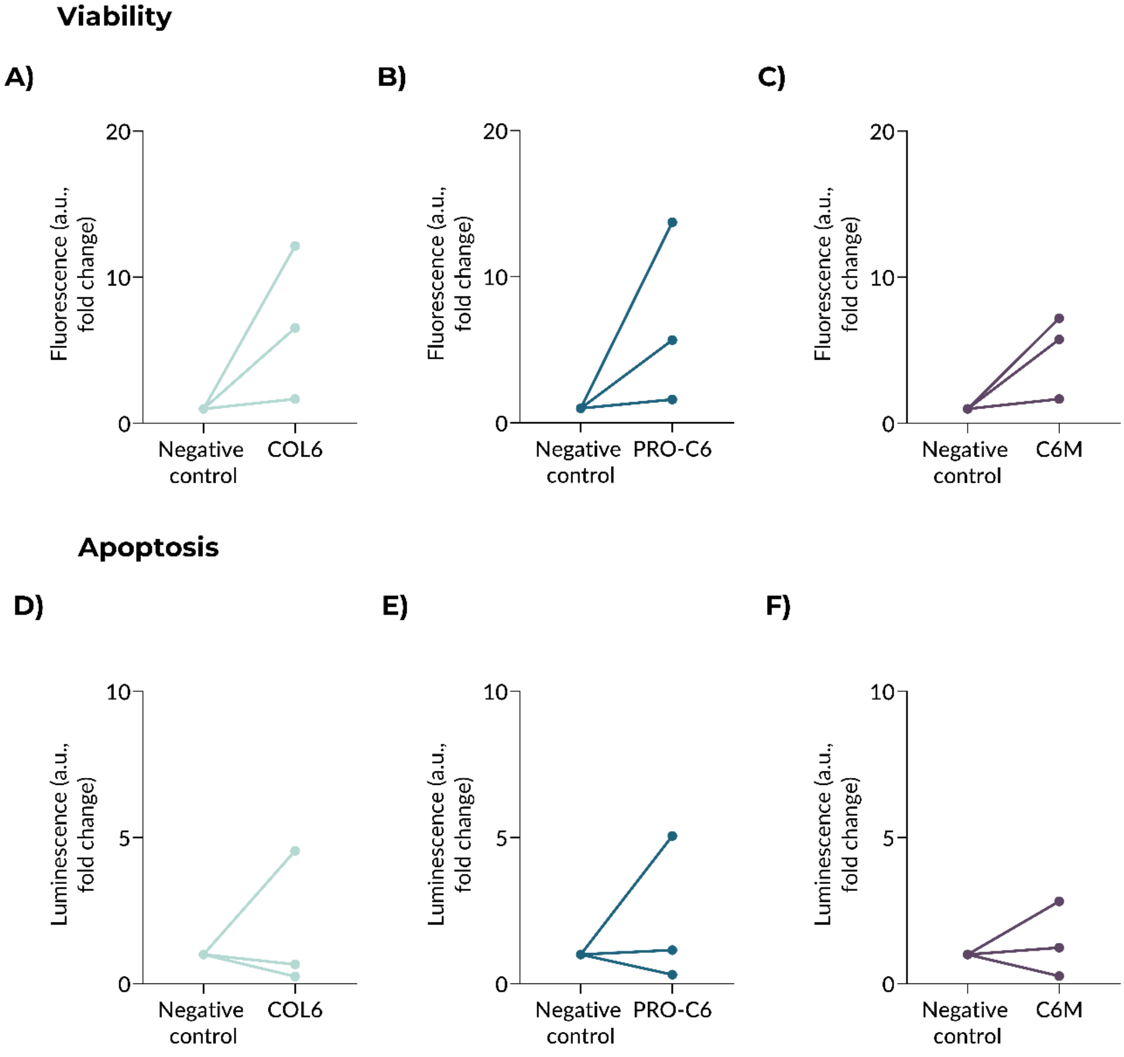


***Supplementary Figure 7**. **COL6, PRO-C6, and C6M tended to increase epithelial cell viability.*** *Brushed epithelial cells were cultured without (negative control) or with 8 ng/µL of COL6, PRO-C6, or C6M for 24h and cell viability and apoptosis were measured using ApoLive-Glo Multiplex Assay. The number of viable cells (viability) shown as fluorescence for the groups: **A)** COL6, **B)** PRO-C6, and **C)** C6M. Caspase activity 3/7 (apoptosis) shown as luminescence for the groups: **D)** COL6, **E)** PRO-C6, and **F)** C6M. Each datapoint represents one biological donor (n=5). Log-transformed data were analyzed with a paired t-test and presented as fold change from the negative control. a.u.: arbitrary unit, C6M: type VI collagen degradation, COL6: type VI collagen, PRO-C6: type VI collagen production.*

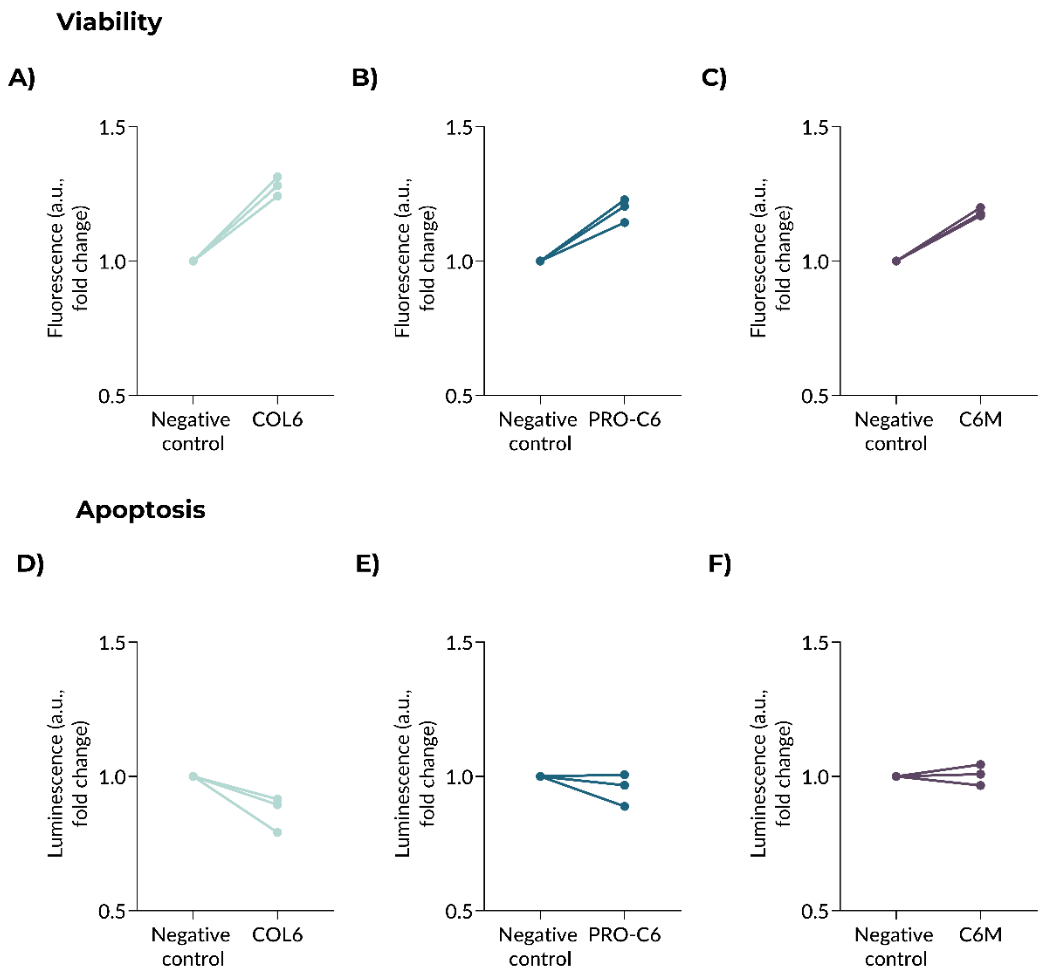


***Supplementary Figure 8*. *COL6, PRO-C6, and C6M tended to increase the viability and decrease the apoptosis of endothelial cells.*** *Human pulmonary micro endothelial cells were cultured without (negative control) or with 8 ng/µL of COL6, PRO-C6, or C6M for 24h and cell viability and apoptosis were measured using ApoLive-Glo Multiplex Assay. The number of viable cells (viability) shown as fluorescence for the groups:* ***A)*** *COL6,* ***B)*** *PRO-C6, and* ***C)*** *C6M. Caspase activity 3/7 (apoptosis) shown as luminescence for the groups:* ***D)*** *COL6,* ***E)*** *PRO-C6, and* ***F)*** *C6M. Each datapoint represents one biological donor (n=5). Log-transformed data were analyzed with a paired t-test and presented as fold change from the negative control. a.u.: arbitrary unit, C6M: type VI collagen degradation, COL6: type VI collagen, PRO-C6: type VI collagen production.*
